# Oscillatory tracking of pseudo-rhythmic speech is constrained by linguistic predictions

**DOI:** 10.1101/2020.12.07.414425

**Authors:** Sanne Ten Oever, Andrea E. Martin

## Abstract

Neuronal oscillations putatively track speech in order to optimize sensory processing. However, it is unclear how isochronous brain oscillations can track pseudo-rhythmic speech input. Here we propose that oscillations can track pseudo-rhythmic speech when considering that speech time is dependent on predictions flowing from internal language models. We show that the temporal dynamics of speech are dependent on the predictability of words in a sentence. A computational model including oscillations, feedback, and inhibition is able to track the natural pseudo-rhythmic speech input. As the model processes, it generates temporal phase codes, which are a candidate mechanism for carrying information forward in time. The model is optimally sensitive to the natural temporal speech dynamics and can explain empirical data on temporal speech illusions. Our results reveal that speech tracking does not only rely on the input acoustics but instead entails an interaction between oscillations and constraints flowing from internal language models.

## Introduction

Speech is a biological signal that is characterized by a plethora of temporal information. The temporal relation between subsequent speech units allows for the online tracking of speech in order to optimize processing at relevant moments in time [1-7]. Neural oscillations are a putative index of such tracking [3, 8]. The existing evidence for neural tracking of the speech envelope is consistent with such a functional interpretation [9, 10]. In these accounts, the most excitable optimal phase of an oscillation is aligned with the most informative time-point within a rhythmic input stream [8, 11-14]. However, the range of onset time difference between speech units seems more variable than fixed oscillations can account for [15-17]. As such, it remains an open question how is it possible that oscillations can track a signal that is at best only pseudo-rhythmic [16].

Oscillatory accounts tend to focus on the prediction in the sense of predicting “when,” rather than predicting “what”: oscillations function to align the optimal moment of processing given that timing is predictable in a rhythmic input structure. If rhythmicity in the input stream is violated, oscillations must be modulated to retain optimal alignment to incoming information. This can be achieved through phase resets [15, 18], directly coupling of the acoustics to oscillations [19], or the use of many oscillators at different frequencies [2]. However, the optimal or effective time of processing stimulus input might not only depend on when you predict something to occur, but also on what stimulus is actually being processed [20-23].

What and when are not independent, and certainly not from the brain’s-eye-view. If continuous input arrives to a node in an oscillatory network, the exact phase at which this node reaches threshold activation does not only depend on the strength of the input, but also on how sensitive this node was to start with. Sensitivity of a node in a language network (or any neural network) is naturally affected by predictions in the what domain generated by an internal language model [24-27]. If a node represents a speech unit that is likely to be spoken next, it will be more sensitive and therefore active earlier, that is, on a less excitable phase of the oscillation. In the domain of working memory, this type of phase precession has been shown in rat hippocampus [28, 29] and more recently in human electroencephalography [30]. In speech, phase of activation and perceived content are also associated [31-35] and phase has been implicated in tracking of higher-level linguistic structure [18, 36, 37]. However, the direct link between phase and the predictability flowing from a language model has yet to be established.

The time of speaking/speed of processing is not only a consequence of how predictable a speech unit is within a stream, but also a cue for the interpretation of this unit. For example, phoneme categorization depends on timing (e.g., voice onsets, difference between voiced and unvoiced phonemes), and there are timing constraints on syllable durations (e.g. the theta syllable [19, 38] that affect intelligibility [39]. Even the delay between mouth movements and speech audio can influence syllabic categorizations [20]. Most oscillatory models use oscillations for parsing, but not as a temporal code for content [40-43]. However, the time or phase of presentation does influence content perception. This is evident from two temporal speech phenomena. In the first phenomena, the interpretation of an ambiguous short /α/ or long vowel /a:/ depends on speech rate (in Dutch; [44-46]). Specifically, when speech rates are fast the stimulus is interpreted as a long vowel and vice versa for slow rates. However, modulating the entrainment rate effectively changes the phase at which the target stimulus - which is presented at a constant speech rate – arrives (but this could not be confirmed in [47]). A second speech phenomena shows the direct phase-dependency of content [31, 34]. Ambiguous /da/-/ga/ stimuli will be interpreted as a /da/ on one phase and a /ga/ on another phase. This was confirmed in both a EEG as well as a behavioral study. An oscillatory theory on speech tracking should account for how temporal properties in the input stream can alter what is perceived.

In the speech production literature, there is strong evidence that the onset times (as well as duration) of an uttered word is modulated by the frequency of that word in the language [48-52] showing that internal language models modulate the access to or sensitivity of a word node [24, 53]. This word-frequency effect relates to the access to a single word. However, it is likely that during ongoing speech internal language models use the full context to estimate upcoming words [54]. If so, the predictability of a word in context should provide additional modulations on speech time. Therefore, we predict that words with a high predictability in the producer’s language model should be uttered relatively early. In this way word-to-word onset times map to the predictability level of that word within the internal model. Thus, not only the processing time depends on the predictability of a word (faster processing for predictable words; see [55, 56] and [57] showing that speech time in noise matters), but also the production time (earlier uttering of predicted words).

Language comprehension involves the mapping of speech units from a producer’s internal model to the speech units of the receiver’s internal model. In other words, one will only understand what someone else is writing or saying if one’s language model is sufficiently similar to the speakers (and if we speak in Dutch, fewer people will understand us). If the producer’s and receiver’s internal language model have roughly matching top-down constrains they should similarly influence the speed of processing (either in production or perception; Figure 1A-C). Therefore, if predictable words arrive earlier (due to high predictability in the producer’s internal model), the receiver also expects the content of this word to match one of the more predictable ones from their own internal model (Figure 1C). Thus, the phase of arrival depends on the internal model of the producer and the expected phase of arrival depends on the internal model of the receiver (Figure 1D). If this is true, pseudo-rhythmicity is fully natural to the brain and it provides a means to use time or arrival phase as a content indicator. It also allows the receiver to be sensitive to less predictable words when they arrive relatively late. Current oscillatory models of speech parsing do not integrate the constraints flowing from an internal linguistic model into the temporal structure of the brain response. It is therefore an open question whether the oscillatory model the brain employs is actually attuned to the temporal variations in natural speech.

**Figure 1.**
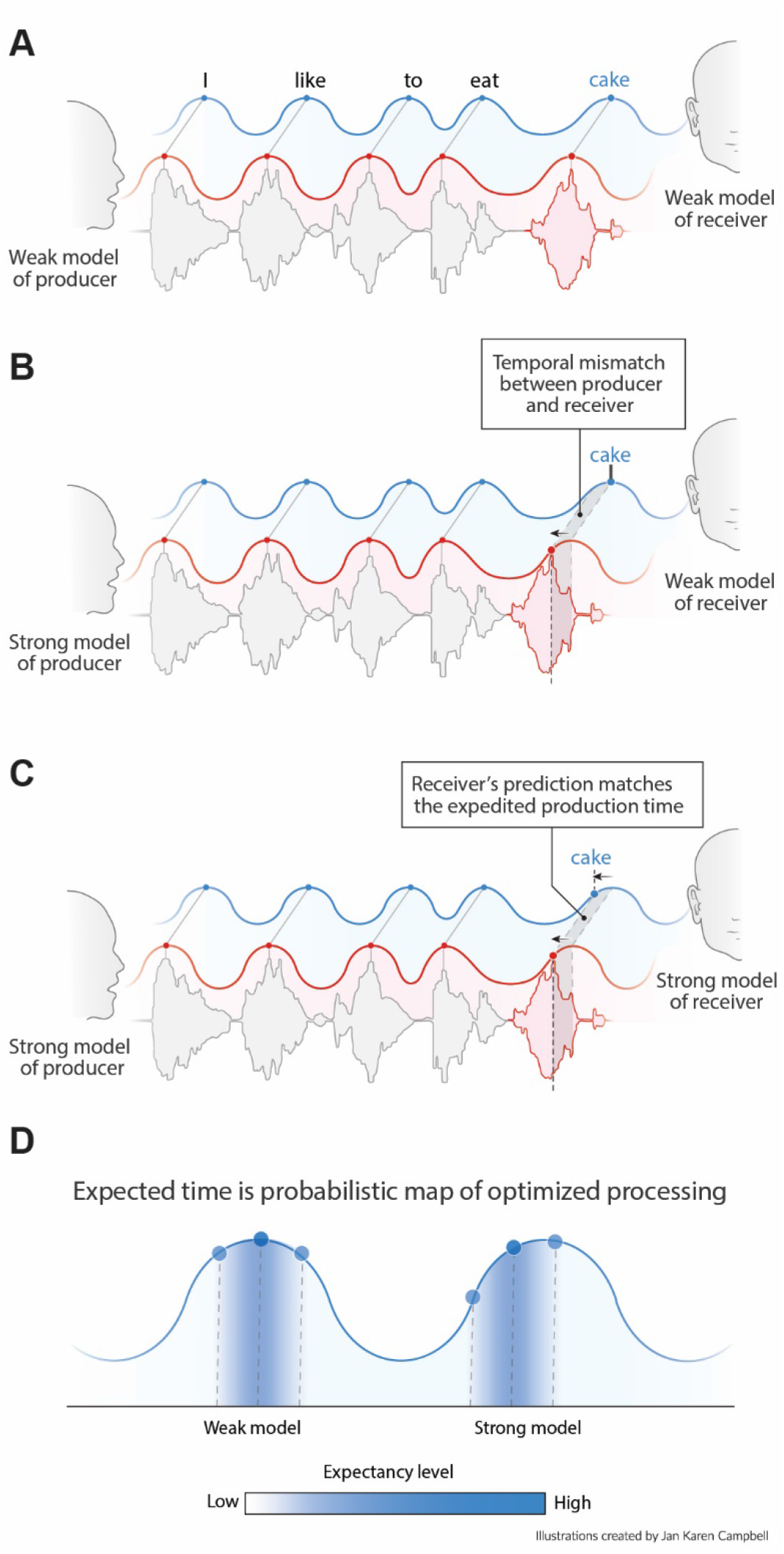
Proposed interaction between speech timing and internal linguistic models. A) Isochronous production and expectation when there is a weak internal model (even distribution of node activation). All speech units arrive around the most excitable phase B) When the internal model of the producer does not align with the model of the receiver temporal alignment and optimal communication fails. C) When both producer and receiver have a strong internal model, speech is non-isochronous and not aligned to the most excitable phase, but fully expected by the brain. D) Expected time is a constraint distribution which center can be shifted due to linguistic constraints.

Here, we propose that neural oscillations can track pseudo-rhythmic speech by taking into account that speech timing is a function of linguistic constrains. As such we need to demonstrate that speech statistics are influenced by linguistic constrains as well as showing how oscillations can be sensitive to this property in speech. We approach this hypothesis as follows: First, we demonstrate that in natural speech timing depends on linguistics predictions (*temporal speech properties*). Then, we model how oscillations can be sensitive to these linguistic predictions (*modeling speech tracking*). Finally, we validate that this model is optimally sensitive to the natural temporal properties in speech and displays temporal speech illusions (*model validation*). Our results reveal that tracking of speech needs to be viewed as an interaction between ongoing oscillations as well as constraints flowing from an internal language model [21, 24]. In this way, oscillations do not have to shift their phase after every speech unit and can remain at a relatively stable frequency as long as the internal model of the speaker matches the internal model of the perceiver.

## Results

### Temporal speech properties: word frequency influences word duration

To extract the temporal properties in naturally spoken speech we used the Corpus Gesproken Nederlands (CGN; (Version 2.0.3; 2014)). This corpus consists of elaborated annotations of over 900 hours of spoken Dutch and Flemish words. We focus here on the subset of the data of which onset and offset timings were manually annotated at the word level in Dutch. Cleaning of the data included removing all dashes and backslashes. Only words were included that were part of a Dutch word2vec embedding (github.com/coosto/dutch-word-embeddings; needed for later modeling) and required to have a frequency of at least 10 in the corpus. All other words were replaced with an <unknown> label. This resulted in 574,726 annotated words with 3096 unique words. 2848 of the words were recognized in the Dutch Wordforms database in CELEX (Version 3.1) in order to extract the word frequency as well as the number of syllables per word. Mean word duration was 0.392 seconds, with an average standard deviation of 0.094 seconds (Supporting Figure 1A). By splitting up the data in sequences of 10 sequential words we could extract the average word, syllable, and character rate (Figure Supporting Figure 1B). The reported rates fall within the generally reported ranges for syllables (5.2 Hz) and words (3.7 Hz; [5, 58]).

We predict that knowledge about the language statistics influences the duration of speech units. As such we predict that more prevalent words will have on average a shorter duration (also reported in [50]). In Figure 2A the duration of several mono- and bi-syllabic words are listed with their word frequency. From these examples it seems that words with higher word frequency generally have a shorter duration. To test this statistically we entered word frequency in an ordinary least square regression with number of syllables as control. Both number of syllables (coefficient = 0.1008, t(2843) = 75.47, p < 0.001) as well as word frequency (coefficient = -0.022, t(2843) = -13.94, p < 0.001) significantly influence the duration of the word. Adding an interaction term did not significantly improve the model (F (1,2843) = 1.320, p = 0.251; Figure 2B+C). The effect is so strong that words with a low frequency can last three times as long as high frequency words (even within mono-syllabic words). This indicates that word frequency could be an important part of an internal model that influences word duration.

**Figure 2.**
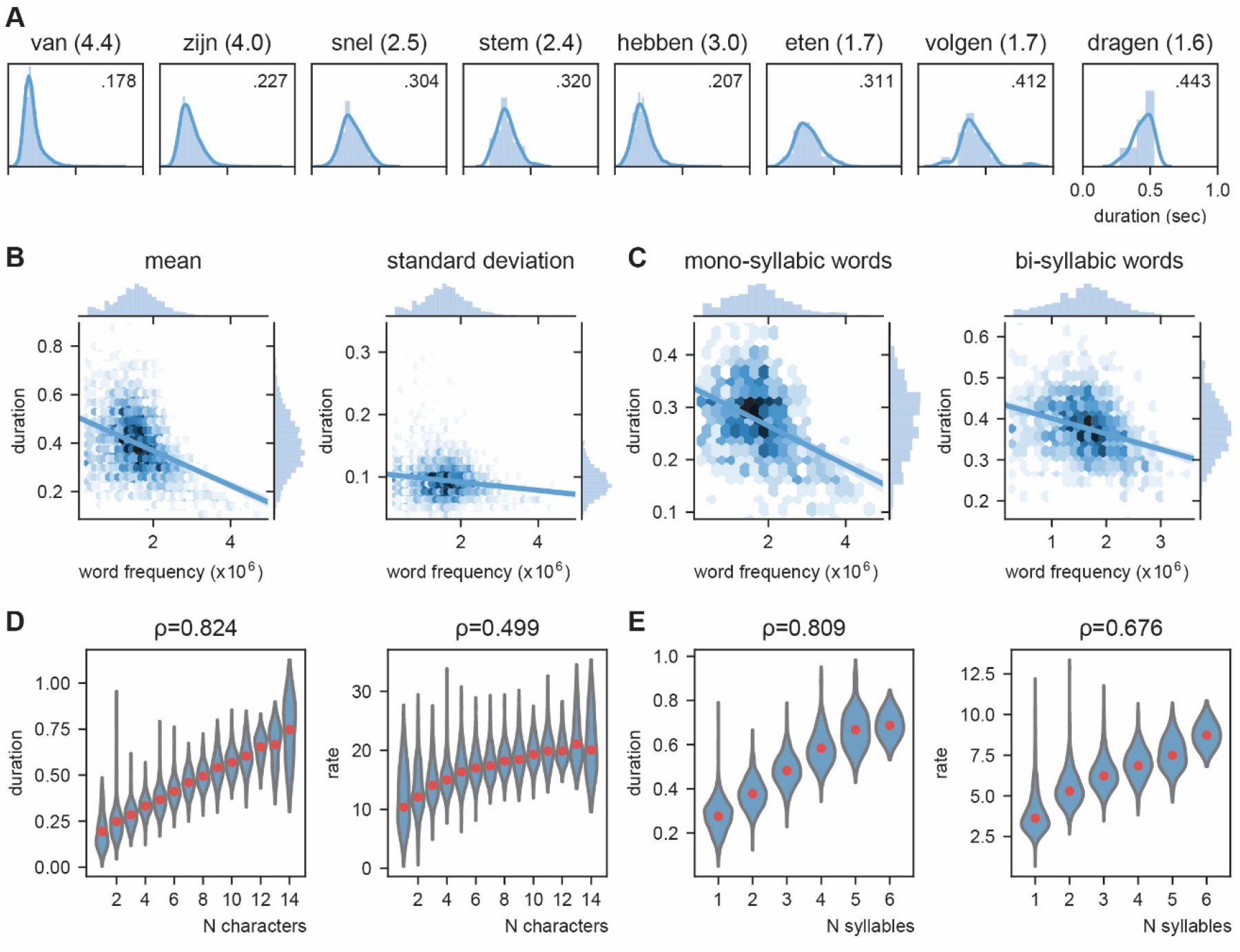
Word frequency modulates word duration. A) Example of mono- and bi-syllabic words of different word frequencies in brackets (van=from, zijn=be, snel=fast, stem=voice, hebben=have, eten=eating, volgend=next, toekomst=future). Text in the graph indicates the mean word duration. B) Relation between word frequency and duration. Darker colors mean more values. C) same as B) but separately for mono- and bi-syllabic words. D) Relation character amount and word duration. The longer the words, the longer the duration (left). The increase in word duration does not follow a fixed number per character as duration as measured by rate increases. E) same as D) but for number of syllables. Red dots indicate the mean.

The previous analysis probed us to expand on the relation between word duration and length of the words. Obviously, there is a strong correlation between word length and mean word duration (number of characters 0.824, p < 0.001; number of syllables: ρ = 0.808, p < 0.001; for number of syllables already shown above; Figure2D+E). In contrast, this correlation is present, but much lower for the standard deviation of word duration (number of characters: ρ = 0.269, p < 0.001; number of syllables: ρ = 0.292, p < 0.001). Finding a strong correlation does not imply that for every time unit increase in the word length, the duration of the word also increases with the same time unit, i.e., bi-syllabic words do not necessarily have to last twice as long as mono-syllabic words. Therefore, we recalculated word duration to a rate unit considering the number of syllables/ characters of the word. Thus a 250 ms mono-versus bi-syllabic word would have a rate of 4 versus 8 Hz respectively. Then we correlated character/syllabic rate with word duration. If word duration increases monotonically with character/syllable length there should be no correlation. We found that the syllabic rate varies between 3 and 8 Hz as previously reported (Figure 2E right; [5, 58]). However, the more syllables there are in a word, the higher this rate (ρ = 0.676, p < 0.001). This increase was less strong for the character rate (ρ = 0.499, p < 0.001; Figure 2D right).

These results show that the syllabic/character rate depends on the number of characters /syllables within a word and is not an independent temporal unit [38]. This effect is easy to explain when assuming that the prediction strength of an internal model influences word duration: transitional probabilities of syllables are simply more constrained within a word than across words [59]. This will reduce the time it takes to utter/perceive any syllable which is later in a word. Unfortunately, the CGN does not have separate syllable annotations to investigate this possibility directly. However, we can investigate the effect of transitional probabilities and other statistical regularities flowing from internal models across words (see next section and [17] for statistical regularities in syllabic processing).

### Temporal speech properties: word-by-word predictability predicts word onset differences

The brain’s internal model likely provides predictions about what linguistic features and representations, and possibly about which specific units, such as words, to expect next when listening to ongoing speech [21, 24]. As such, it is also expected that word-by-word onset delays are shorter for words that fit the internal model (i.e. those that are expected; [54]). To investigate this possibility, we created a simplified version of an internal model predicting the next word using recurrent neural nets (RNN). We trained an RNN to predict the next word from ongoing sentences (Figure 3A). The model consisted of an embedding layer (pretrained; github.com/coosto/dutch-word-embeddings), a recurrent layer with a tanh activation function, and a dense output layer with a softmax activation. To prevent overfitting, we added a 0.2 dropout to the recurrent layers and the output layer. An adam optimizer was used at a 0.001 learning rate and a batch size of 32. We investigated four different recurrent layers (GRU and LSTM at either 128 or 300 units; see Supporting Figure 4). The final model we use here includes a LSTM with 300 units. Input data consistent of 10 sequential words (label encoding) within the corpus (of a single speaker; shifting the sentences by one word at a time), and an output consisted of a single word. A maximum of four unknown labeled words (words not included in the word2vec estimations. Four was choosen as it was < 50% of the words). was allowed in the input, but not in output. Validation consisted of a randomly chosen 2% of the data.

**Figure 3.**
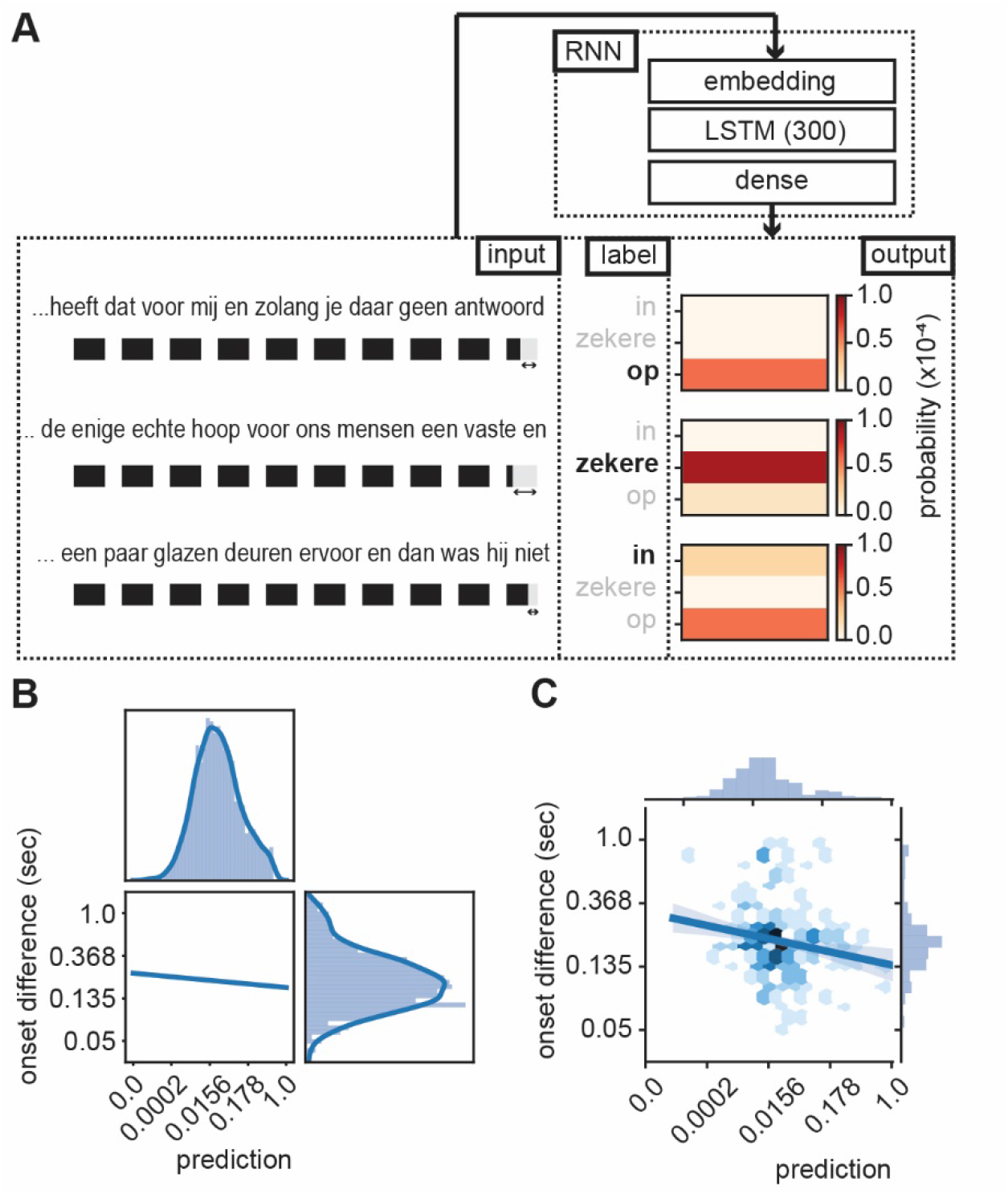
RNN output influence word onset differences. A) Sequences of ten words were entered in an RNN in order to predict the content of the next word. Three examples are provided of input data with the label (bold word) and probability output for three different words. The regression model showed a relation between the duration of last word in the sequence and the predictability of the next word such that words were systematically shorter when the next word was more predictable according to the RNN output (illustrated here with the shorted black boxes). B) Regression line estimated at mean value of word duration and bigram. C) Scatterplot of prediction and onset difference of data within ± 0.5 standard deviation of word duration and bigram. Note that for B and C the axes are linear on the transformed values. Translation of the sentences in A from top to bottom: *“*… *that it has for me and while you have no answer [on]”, “*… *the only real hope for us humans is a firm and [sure]”, “*… *a couple of glass doors in front and then it would not have been [in]”*.

The output of the RNN reflects a probability distribution in which the values of the RNN sum up to one and each word has its own predicted value (Figure 3A). As such we can extract the predicted value of the uttered word and relate the RNN prediction with the stimulus onset delay relative to the previous word. We entered word prediction in a regression using the stimulus onset difference between the current word in the sentence and the previous word (i.e. onset difference of words). We added the control variables bigram (using the NLTK toolbox based on the training data only), frequency of previous word, syllable rate (rate of the full sentence input), and mean duration of previous word (all variables that can account for part of the variance that affects the duration of the last word). We only used the test data (total of 7361 sentences, excluding all word not present in Celex. 4837 sentences). Many of the variables were skewed to the right, therefore we transformed the data accordingly (see Table 1; results were robust to changes in these transformation).

**Table 1.** Summary of regression model for logarithm of onset difference of words

| Variable | Trans | B | $\beta$ | SE | t | p | VIF |
| --- | --- | --- | --- | --- | --- | --- | --- |
| Intercept | x | 0.9719 |  | 0.049 | 19.764 | <0.001 |  |
| RNN prediction | $x^{(1/6)}$ | -0.3370 | -0.0862 | 0.047 | -7.163 | <0.001 | 1.5 |
| Bigram | $\log(x)$ | -0.0118 | -0.0316 | 0.005 | -2.424 | 0.015 | 1.8 |
| Word frequency W-1 | x | 0.0049 | 0.0076 | 0.009 | 0.546 | 0.585 | 2.0 |
| Mean duration W-1 | $\log(x)$ | 1.1206 | 0.7003 | 0.022 | 50.326 | <0.001 | 2.0 |
| Syllable Rate | x | -0.1033 | -0.2245 | 0.004 | -23.014 | <0.001 | 1.0 |

Model $R^2 = 0.542$ . Trans = transformation, W-1 = previous word, B = unstandardized coefficient, $\beta$ = standardized coefficient, SE = standard error, t = t value, p = p value, VIF = variance inflation factor

All predictors except word frequency of the previous word showed a significant effect (Table 1). The variance explained by word frequency was likely captured by the mean duration variable of the previous word which correlated word frequency. The RNN predictor could capture more variance than the bigram model suggesting that word duration is modulated by the level of predictability within a fuller context than just the conditional probability of the current word given the previous word (Figure 3B+C). Importantly, it was necessary to use the trained RNN model as a predictor; entering the RNN predictions after the first epoch did not results in a significant predictor (t(4837) = -1.191, p = 0.234). Also adding the predictor word frequency of the current word did not add significant information to the model (F(1, 4830) = 0.2048, p = 0.651). These results suggest that words are systematically lengthened (or pauses are added. However, the same predictors are also significant when excluding sentences containing pauses) when the next word is not strongly predicted by the internal model.

### Modeling speech tracking: Speech Tracking in a Model Constrained Oscillatory Network (STiMCON)

In order to investigate how much of these duration effects can be explained using an oscillator model, we created the model Speech Tracking in a Model Constrained Oscillatory Network (STiMCON). STiMCON in its current form will not be exhaustive; however, it can extract how much an oscillating network can cope with asynchronies by using its own internal model illustrating how the brain’s language model and speech timing interact [60]. The current model is capable of explaining how top-down predictions can influence the processing time as well as provide an explanation for two known temporal illusions in speech.

STiMCON consists of a network of semantic nodes of which the activation A of each level in the model l is governed by:

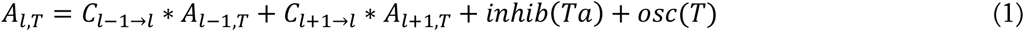

in which C represents the connectivity patterns between differrent hierarchical levels, T the time in a sentence, and Ta the vector of times of an individual node in an inhibition function (in milliseconds). The inhibition function is a gate function:

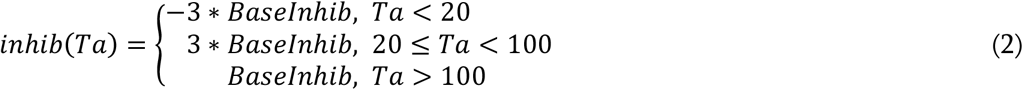

in which BaseInhib is a constant for the base level of inhibition (negative value, set to -0.2). As such nodes are by default inhibited, as soon as they get activated above threshold (activation threshold set at 1) Ta sets to zero. Then, the node will have suprathreshold activation, which after 20 milliseconds returns to increased inhibition until the base level of inhibition is returned. The oscillation is a constant oscillator:

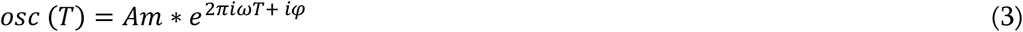

in which Am is the amplitude of the oscillator, ω the frequency, and φ the phase offset. As such we assume a stable oscillator which is already aligned to the average speech rate (see [15, 19] for phase alignment models). The model used for the current simulation has one an input layer (l-1 level) and one single layer of semantic word nodes (l level) that receives feedback from a higher level layer (l+1 level). As such only the word (l) level is modeled according to equation 1-3 and the other levels form fixed input and feedback connection patterns.

### Modeling speech tracking: language models influence time of activation

To illustrate how STiMCON can explain how processing time depends on the prediction of internal language models, we instantiated a language model that had only seen three sentences and five words presented at different probabilities (I eat cake at 0.5 probability, I eat nice cake at 0.3 probability, I eat very nice cake at 0.2 probability; Table 2). This language model will serve as the feedback arriving from the l+1-level to the l-level. The l-level consists of five nodes that each represent one of the words and receives proportional feedback from l+1 according to Table 2 with a delay of 0.9*ω seconds, which then decays at 0.01 unit per millisecond and influences the l-level at a proportion of 1.5. This feedback is only initiated when supra-activation arrives due to l-1-level bottom-up input. Each word at the l-1-level input is modelled as a linearly function to the individual nodes lasting length of 125 milliseconds (half a cycle, ranging from 0-1 arbitrary units). As such, the input is not the acoustic input itself but rather reflects a linear increase representing the increasing confidence of a word representing the specific node. φ is set such that the peak of a 4 Hz oscillation aligns to the peak of sensory input of the first word. Sensory input is presented at a base stimulus onset asynchrony of 250 milliseconds (i.e. 4 Hz).

**Table 2.** Example of a language model.

|  | I | eat | very | nice | cake |
| --- | --- | --- | --- | --- | --- |
| I | 0 | 1 | 0 | 0 | 0 |
| eat | 0 | 0 | 0.2 | 0.3 | 0.5 |
| very | 0 | 0 | 0 | 1 | 0 |
| nice | 0 | 0 | 0 | 0 | 1 |
| cake | 0 | 0 | 0 | 0 | 0 |

This model has seen three sentences at different probabilities. Rows represent the prediction for the next word, e.g. /I/ predicts /eat/ at a probability of 1, but after /eat/ there is a wider distribution.

When we present this model with different sensory input at an isochronous rhythm of 4 Hz it is evident that the timing at which different nodes reach activation depends on the level of feedback that is provided (Figure 4). For example, while the /I/-node needs a while to get activated after the initial sensory input, the /eat/-node is activated earlier as it is pre-activated due to feedback. After presenting /eat/ the feedback arrives at three different nodes and the activation timing depends on the stimulus that is presented (earlier activation for /cake/ compared to /very/).

**Figure 4.**
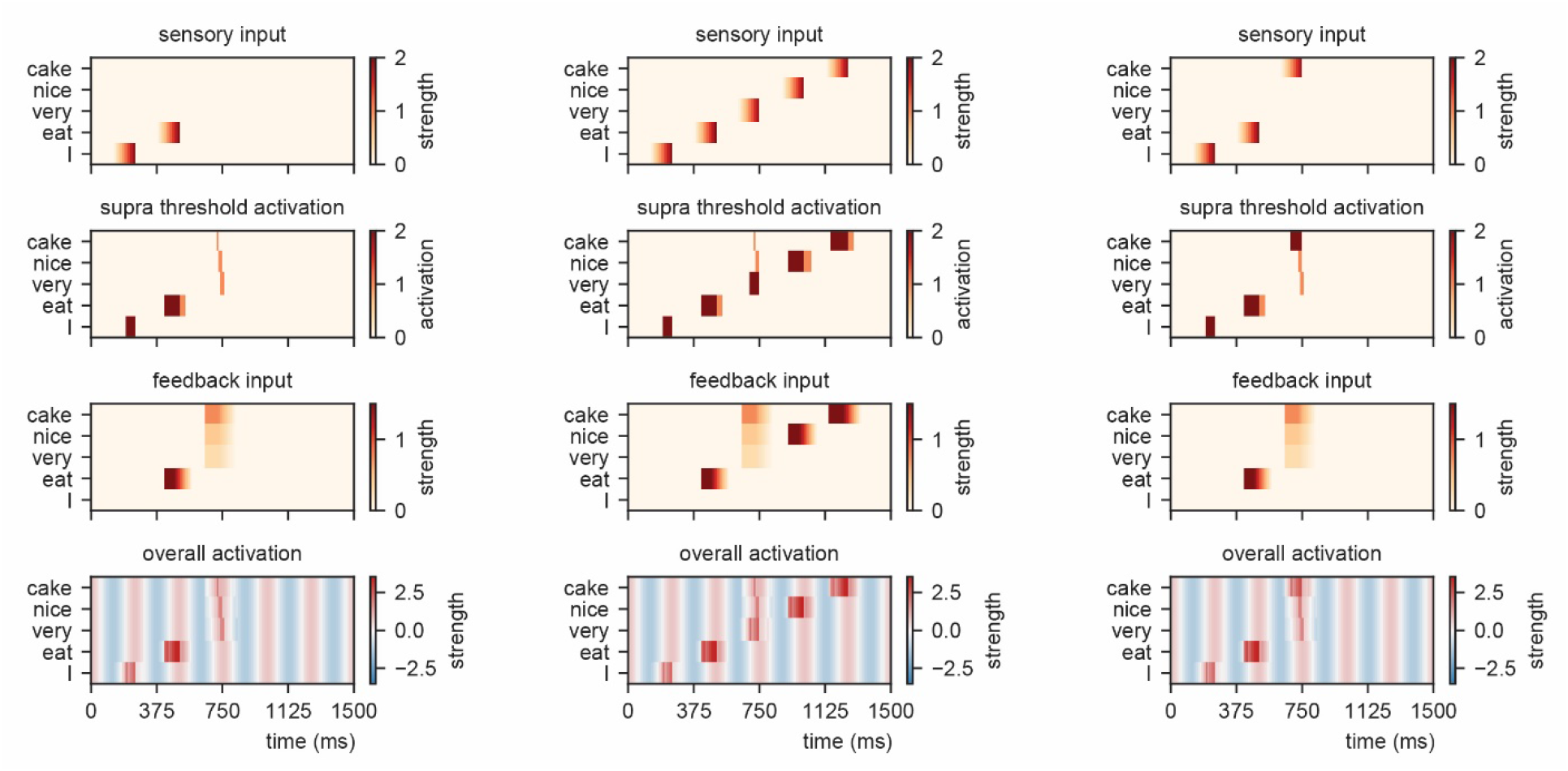
Model output for different sentences. For the supra-threshold activation dark red indicates activation which included input from *l+1* as well as *l-1*, orange indicates activation due to *l+1* input.

### Modeling speech tracking: time of presentation influences processing efficiency

To investigate how the time of presentation influences the processing efficiency we presented the model with /I eat XXX/ in which the last word was varied in content (either /I/, /very/, /nice/, or /cake/), intensity (linearly ranging from 0 to 1), and onset delay (ranging between -125 to +125 relative to isochronous presentation). We extracted the time at which the node matching the stimulus presentation reached activation threshold first (relative to stimulus onset, and relative to isochronous presentation).

Figure 5A shows the output. When there is no feedback (i.e. at the first word /I/ presentation), a classical efficiency map can be found in which processing is most optimal (possible at lowest stimulus intensities) at isochronous (in phase with the stimulus rate) presentation and then drops to either side. For nodes that have feedback, input processing is possible at earlier times relative to isochronous presentation and parametrically varies with prediction strength (earlier for /cake/ at 0.5 probability, then /very/ at 0.2 probability). Additionally, the activation function is asymmetric. This is a consequence of the interaction between the supra-activation caused by the feedback and the sensory input. As soon as supra-activation is reached due to the feedback, sensory input at any intensity will reach supra-activity (thus at early stages of the linearly increasing confidence of the input). This is why for the /very/ stimulus activation is still reached at later delays compared to /nice/ and /cake/ as the /very/-node reaches supra-activation due to feedback at a later time point.

**Figure 5.**
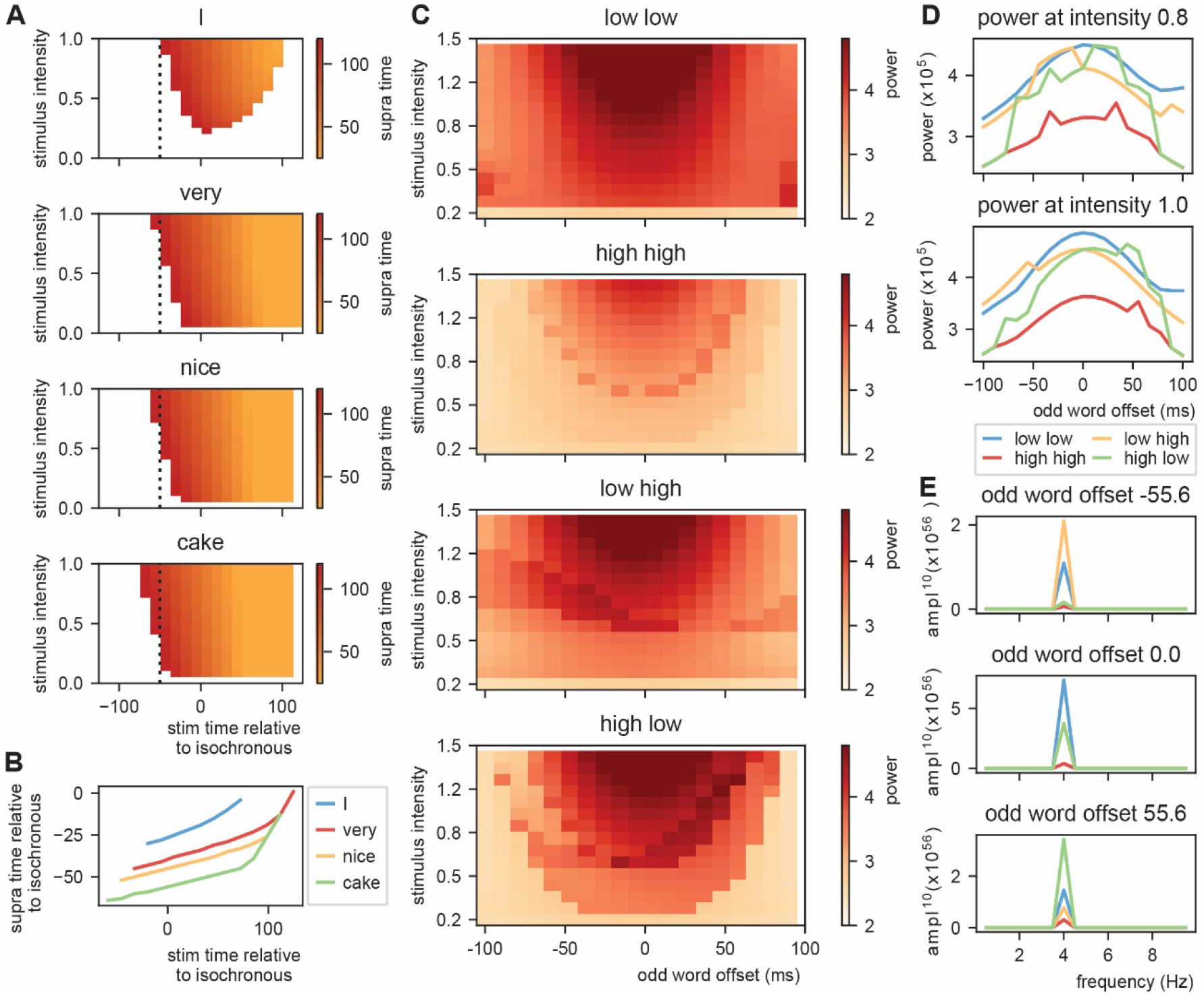
Model output on processing efficiency and rhythmicity. A) Time of presentation influences efficiency. Outcome variable is the time at which the node reached threshold activation (supra-time). The dashed line is presented to ease comparison between the four content types. White indicates that threshold is never reached. B) Same as A, but estimated at a threshold of 0.53 showing that oscillations regulate feedforward timing. Panel A shows that the earlier the stimuli are presented (on a weaker point of the ongoing oscillation), the longer it takes until supra-threshold activation is reached. This figure shows that timing relative to the ongoing oscillation is regulated such that the stimulus activation timing is closer to isochronous. Line discontinuities are a consequence of stimuli never reaching threshold for a specific node. C) Strength of 4 Hz power depends on predictability in the stream. When predictability is alternated between low and high, activation is more rhythmic when the predictable odd stimulus arrives earlier and vice versa. D) Slice of D at intensity of 0.8 and 1.0. E) Magnitude spectra at three different odd word offsets at 1.0 intensity. To more clearly illustrate the differences the magnitude to the power of 20 is plotted.

When we investigate timing differences in stimulus presentation it is important to also consider what this means for the timing in the brain. Before, we showed that the amount of prediction can influence timing in our model. It is also evident that the earlier a stimulus was presented the more time it took (relative to the stimulus) for the nodes to reach threshold (more yellow colors for earlier delays). This is a consequence of the oscillation still being at a relatively low excitability point at stimulus onset for stimuli that are presented early during the cycle. However, when we translate these activation threshold timing to the timing of the ongoing oscillation, the variation is strongly reduced (Figure 5B). A stimulus timing that varies between 130 milliseconds (e.g. from -59 to +72 in the /cake/ line; excluding the non-linear section of the line) only reaches the first supra-threshold response with 19 milliseconds variation in the model (translating to a reduction of 53% to 8% of the cycle of the ongoing oscillation, i.e. a 1:6.9 ratio). This means that within this model (and any oscillating model) the activation of nodes is robust to some timing variation in the environment. This effect seemed weaker when no prediction was present (for the /I/ stimulus this ratio was around 1:3.5. Note that when determining the /cake/ range using the full line the ratio would be 1:3.4).

### Modeling speech tracking: top-down interactions can provide rhythmic processing for non-isochronous stimulus input

The previous simulation demonstrate that oscillations provide a temporal filter and the processing itself can actually be closer to isochronous than what can be solely extracted from the stimulus input. Next, we investigated whether dependent on changes in top-down prediction, processing within the model will be more or less rhythmic. To do this, we create stimulus input of 10 sequential words at a base rate of 4 Hz to the model with constant (low at 0 and high at 0.8 predictability) or alternating word-to-word predictability. For the alternating conditions word-to-word predictability alternates between low to high (sequences which word are predicted at 0 or 0.8 predictability, respectively) or shift from high to low. For this simulation we used Gaussian sensory input (with a standard deviation of 42 ms aligning the mean at the peak of the ongoing oscillation; see Supporting Figure 5 for output with linear sensory input). Then, we vary the onset time of the odd words in the sequence (shifting from -100 up to +100 ms) and the stimulus intensity (from 0.2 to 1.5). We extracted the overall activity of the model and computed the Fast Fourier transform of the created time course (using a Hanning taper only including data from 0.5 – 2.5 seconds to exclude the onset responses).

The first thing that is evident is that the model with no content predictions has overall stronger power, and specifically around isochronous presentation (odd word offset of 0 ms) at high stimulus intensities (Figure 5C-E). Adding overall high predictability drops the power, but also here the power seems symmetric around zero. The spectra of the alternating predictability conditions look different. For the low to high predictability condition the curve seems to be shifted to the left such that 4 Hz power is strongest when the predictable odd stimulus is shifted to an earlier time point (low-high condition). This is reversed for the high-low condition. At middle stimulus intensities there is a specific temporal specificity window at which the 4 Hz power is particularly strong. This window is earlier for the low-high than the high-low alternation (Figure 5D, Figure 5E, and Supporting Figure 6). The effect only occurs at specific middle intensity combination as at high intensities the stimulus dominates the responses and at low intensities the stimulus does not reach threshold activation. These results show that even though stimulus input is non-isochronous, the interaction with the internal model can still create a potential rhythmic structure in the brain (see [61, 62]). Note that the direction in which the brain response is more rhythmic matches with the natural onset delays in speech (shorter onset delays for more predictable stimuli).

### Model validation: STiMCON’s sinusoidal modulations of RNN predictions is optimally sensitive to natural onset delays

Next, we aimed to investigated whether STiMCON would be optimally sensitive to speech input timings found naturally in speech. Therefore, we tried to fit STIMCON’s expected word-to-word onset differences to the word-to-word onset differences we found in the CGN. At a stable level of intensity of the input and inhibition, the only aspect that changes the timing of the interaction between top-down predictions and bottom-up input within STiMCON is the ongoing oscillation. Considering that we only want to model for individual words how much the prediction (*C*_*l−*1→*l*_ * *A*_*l−*1*,T*_) influences the expected timing we can set the contribution of the other factors from equation (1) to zero remaining with the relative contribution of prediction:

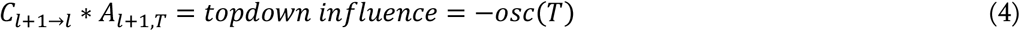

We can solve this formula in order to investigate the expected relative time shift (T) in processing that is a consequence of the strength of the prediction (ignoring that in the exact timing will also depend on the strength of the input and inhibition):

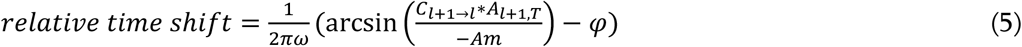

ω was set as the syllable rate for each sentence, Am and φ were systematically varied. We fitted a linear model between the STiMCON’s expected time and the actual word-to-word onset differences. This model was similar to the model described in the section *word-by-word predictability predicts word onset differences* and included the predictor syllablerate and duration of the previous word. However, as we were interested in how well non-transformed data matches the natural onset timings we did not perform any normalization besides equation (5). As this might involve violating some of the assumptions of the ordinary least square fit, we estimate model performance by repeating the regression 1000 times fitting it on 90% of the data (only including the test data from the RNN) and extracting R^2^ from the remaining 10%.

Results show a modulation of the R^2^ dependent on the amplitude and phase offset of the oscillation (Figure 6A) which was stronger than the non-transformed R^2^ (which was 0.389). This suggests that STiMCON expected time durations matches the actual word-by-word duration. This was even more strongly so for specific oscillatory alignments (around -0.25π offset) suggesting an optimal alignment phase relative to the ongoing oscillation is needed for optimal tracking [3, 8]. Interestingly, the optimal transformation seemed to automatically alter a highly skewed prediction distribution (Figure 6B) towards a more normal distribution of relative time shifts (Figure 6C).

**Figure 6.**
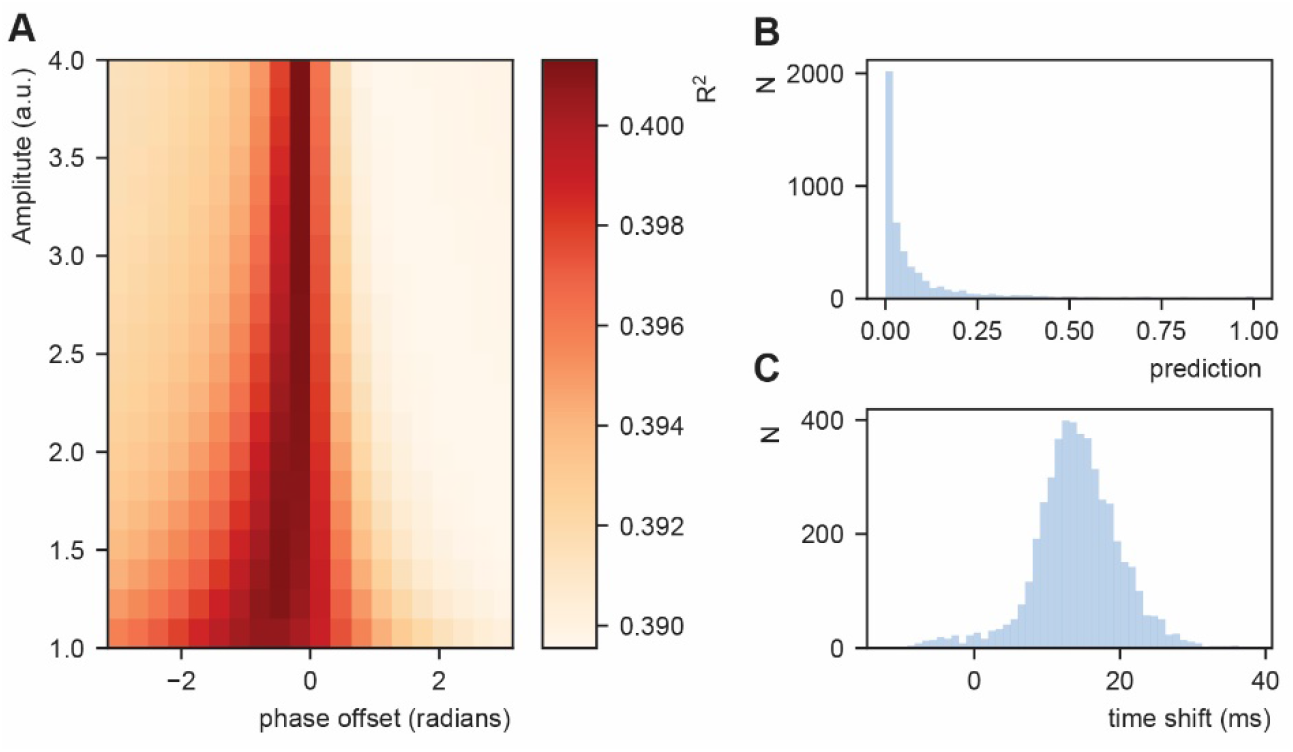
Fit between real and expected time shift dependent on predictability. A) Phase offset and amplitude of the oscillation modulate the fit to the word-to-word onset durations. B) Histogram of the predictions created by the deep neural net. C) Histogram of the relative time shift transformation at phase of -0.15π and amplitude of 1.5.

### Model validation: STiMCON can explain perceptual effects in speech processing

Due to the differential feedback strength and the inhibition after suprathreshold feedback stimulation, STiMCON is more sensitive to lower predictable stimuli at phases later in the oscillatory cycle. This property can explain two illusions that have been reported in the literature, specifically, the observation that the interpretation of ambiguous input depends on the phase of presentation [31, 32, 63] and on speech rate [46]. The only assumption that has to be made is that there is an uneven base prediction balance between the ways the ambiguous stimulus can be interpreted.

The empirical data we aim to model comprises an experiment in which ambiguous syllables, that could either be interpreted as /da/ or /ga/, were presented [31]. In one of the experiments in this study, broadband simuli were presented at specific rates to entrain ongoing oscillations. After the last entrainment stimulus an ambiguous /daga/ stimulus was presented at different delays (covering two cycles of the presentation rate at 12 different steps), putatively reflecting different oscillatory phases. Dependent on the delay of stimulation participants perceived either /da/ or /ga/ suggesting that phase modulates the percept of the participants. Besides this behavioral experiment, the authors also demonstrated that the same temporal dynamics were present when looking at ongoing EEG data showing that the phase of ongoing oscillations at the onset of ambiguous stimulus presentation determined the percept [31].

To illustrate that STiMCON is capable of showing a phase (or delay) dependent effect, we use an internal language model similar to our original model (Table 2). The model consists of four nodes (N1, N2, Nda, and Nga) at which N1 activation predicts a second unspecific stimulus (S2) represented by N2 at a predictability of 1. N2 activation predicts either da or ga at 0.2 and 0.1 probability respectively. Then, we present STiMCON (same parameters as before) with /S1 S2 XXX/. XXX is varied to have different proportion of the stimulus /da/ and /ga/ (ranging from 0% /da/ to 100% /ga/ in 12 times steps; these reflects relative propotions that sum up to 1 such that at 30% the intensity of /da/ would be at max 0.3. and of /ga/ 0.7) and is the onset is varied relate to the second to last word. We extract the time that a node reaches suprathreshold activity after stimulus onset. If both nodes were active at the same time the node with the highest total activates was choosen. Results showed that for some ambiguous stimuli, the delay determines which node is activated first, modulating the ultimate percept of the participant (Figure 7A, also see supplementary Figure 7A). The same type of simulation can explain how speech rate can influence perception (supplementary Figure 7B; but see [47].).

**Figure 7.**
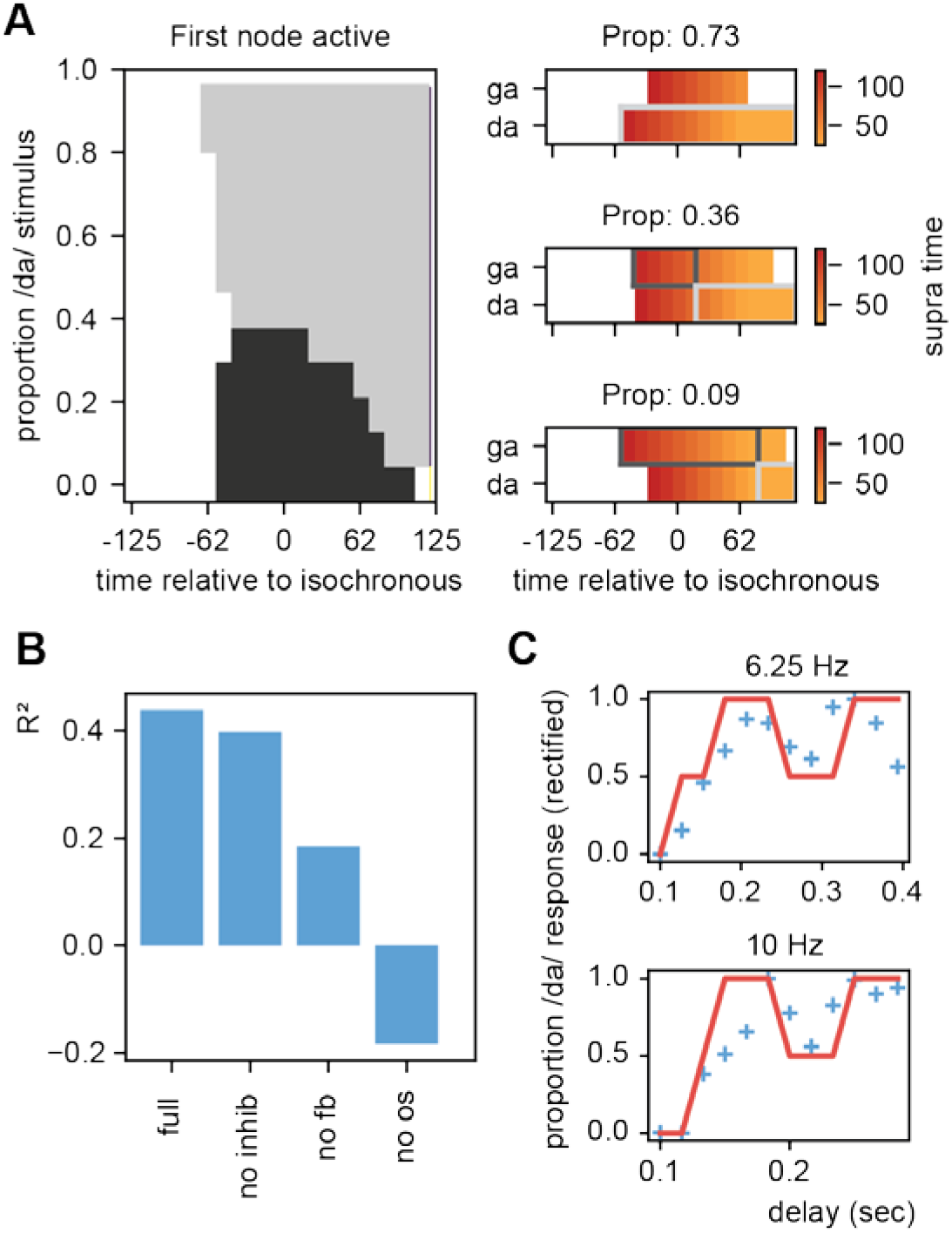
Results for /daga/ illusions. A) Modulations due to ambiguous input at different times. Illustration of the node that is active first. Different proportions of the /da/ stimulus show activation timing modulations at different delays (B). R^2^ for the grid search fit of the full model, a model without inhibition (no inhib), without uneven feedback (no fb), or without an oscillation (no os). C) Fit of the full model on the rectified behavioral data of [31]. Blue crossed indicate rectified data and red lines indicate the fit.

To further scrutinize on this effect we fitted our model to the behavioral data of Ten Oever & Sack [31]. As we used an iterative approach in the simulations of the model, we optimized the model using a grid search. We varied the parameters of proportion of the stimulus being /da/ or /ga/ (ranging between 10:20:80%), the onset time of the feedback (0.1:0.2:1.0 cycle), the speech of the feedback decay (0:0.02:0.1), and a temporal offset of the final sound to account for the time it takes to interpret a specific ambiguous syllable (ranging between -0.05:0.02:0.05 sec). Our outcome variable was the node that show the first suprathreshold activation (Nda = 1, Nga = 0). If both nodes were active at the same time the node with the highest total activates was choosen. If both nodes had equal activation or never reached threshold activation we coded the outcome to 0.5 (i.e. fully ambigous). These outcomes were fitted to the behavioral data of the 6.25 Hz and 10 Hz presentation rate (the two rates showing a significant modulation of the percept). This data was normalize to have a range between 0-1 to account for the model outcomes being binary (0, 0.5 or 1).

We found that our model could fit the data at an average explained variance of 43% (30% and 58% for 6.25 Hz and 10 Hz respectively; Figure 7B+C). This explained variance was higher than the original sinus fit (40% for 3 parameter sinus fit [amplitude, phase offset, and mean]). Note that our fit cannot account for variance ranging inbetween 0-0.5 and 0.5-1, while the sinus fit can do this. If we correct for this (by setting the sinus fit to the closest 0, 0.5 or 1 value and doing a grid search to optimize the fitting) the average fit of the sinus is 21%. The average AIC of the model and sinus fit are -27.0 and -24.1 respectively suggesting that the STiMCON model has the better fit. Thus, STiMCON does better than a fixed-frequency sinus fit. This is a likely consequence of the sinus fit not being able to explain the dampening of the oscillation later (i.e. the perception bias is stronger for shorter compared to longer delays).

Finally, we investigated the relevance of the three key features of our model for this fit: inhibition, feedback, and oscillations. We repeated the grid search fit but set either the inhibition to zero, the feedback matrix equal for both /da/ and /ga/ (both 0.15), or the oscillation at an amplitude of zero. Results showed that especially the oscillation and the differential feedback were essential to reach a good fit (Figure 7B). Without the oscillation the model could not even fit better than the mean of the model (R^2^ < 0). Removing the inhibition had the least influence on the fit. This suggest that all features (with a lesser extend the inhibition) are required to model the data suggesting that oscilatory tracking is dependent on linguistic constrains flowing from the internal language model.

## Discussion

In the current paper, we combined an oscillatory model with a proxy for linguistic knowledge, an internal language model, in order to investigate the model’s processing capacity for onset timing differences in natural speech. We show that word-to-word speech onset differences in natural speech are indeed related to predictions flowing from the internal language model (estimated through an RNN). Fixed oscillations aligned to the mean speech rate are robust against natural temporal variations and even optimized for temporal variations that match the predictions flowing from the internal model. Strikingly, when the pseudo-rhythmicity in speech matches the predictions of the internal model, responses were more rhythmic for matched pseudo-rhythmic compared to isochronous speech input. Our model is optimally sensitive to natural speech variations, can explain phase dependent speech categorization behavior [31, 35, 44, 63], and naturally comprises a neural phase code [40, 42, 43]. These results show that part of the pseudo-rhythmicity of speech is expected by the brain and it is even optimized to process it in this manner, but only when it follows the internal model.

Speech timing is variable and in order to understand how the brain tracks this pseudo-rhythmic signal we need a better understanding of how this variability arises. Here, we isolated one of the components explaining speech time variation, namely, constraints that are posed by an internal language model. This goes beyond extracting the average speech rate [5, 19, 58], and might be key to understanding how a predictive brain uses temporal cues. We show that speech timing depends on the predictions made from an internal language model, even when those predictions are highly reduced to be as simple as word predictability. While syllables generally follow a theta rhythm, there is a systematic increase in syllabic rate as soon as more syllables are in a word. This is likely a consequence of the higher close probability of syllables within a word which reduces the onset differences of the later uttered syllables [59]. However, an oscillatory model constrained by an internal language model is sensitive to these temporal variations, it is actually capable of processing them optimally.

The oscillatory model we here pose has three components: oscillations, feedback, and inhibition. The oscillations allow for the parsing of speech and provide windows in which information is processed [3, 39, 64, 65]. Importantly, the oscillation acts as a temporal filter, such that the activation time of any incoming signal will be confined to the high excitable window and thereby is relatively robust against small temporal variations (Figure 5B). The feedback allows for differential activation time dependent on the sensory input (Figure 5A). As a consequence, the model is more sensitive to higher predictable speech input and therefore active earlier on the duty cycle (this also means that oscillations are less robust against temporal variations when the feedback is very strong). The inhibition allows for the network to be more sensitive to less predictable speech units when they arrive later (the higher predictable nodes get inhibited at some point on the oscillation; best illustrated by the simulation in Figure 7A). However, adding inhibition only slightly improved the modeling fit (Figure 7B). In this way speech is ordered along the duty cycle according to its predictability [43, 66]. The feedback in combination with an oscillatory model can explain speech rate and phase dependent content effects. Moreover, it is an automatic temporal code that can use time of activation as a cue for content [42]. The three components in the model are common brain mechanisms [29, 42, 67-70] and follow many previously proposed organization principles (e.g. temporal coding and parsing of information). While we implement these components on an abstract level (not veridical to the exact parameters of neuronal interactions), they illustrate how oscillations, feedback, and inhibition interact to optimize sensitivity to natural pseudo-rhythmic speech.

The current model is not exhaustive and does not provide a complete explanation of all the details of speech processing in the brain. For example, it is likely that the primary auditory cortex is still mostly modulated by the acoustic pseudo-rhythmic input and only later brain areas follow more closely the constraints posed by the language model of the brain. Therefore, more hierarchical levels need to be added to the current model (but this is possible following equation (1)). Moreover, the current model does not allow for phase or frequency shifts. This was intentional in order to investigate how much a fixed oscillator could explain. We show that onset times matching the predictions from the internal model can be explained by a fixed oscillator processing pseudo-rhythmic input. However, when the internal model and the onset timings do not match the internal model phase and/or frequency shift are still required and need to be incorporated (see e.g. [15, 19]). Still, any coupling between brain oscillations and speech acoustics [19] needs to be extended with the coupling of brain oscillations to brain activity patterns of internal models [71].

In the current paper we use an RNN to represent the internal model of the brain. However, it is unlikely that the RNN captures the wide complexities of the language model in the brain. The decades-long debates about the origin of a language model in the brain remains ongoing and controversial. Utilizing the RNN as a proxy for our internal language model makes a tacit assumption that language is fundamentally statistical or associative in nature, and does not posit the derivation or generation of knowledge of grammar from the input [72, 73]. In contrast, our brain could as well store knowledge of language that functions as fundamental interpretation principles to guide our understanding of language input [21, 24, 53, 65, 74]. Knowledge of language and linguistic structure could be acquired through an internal self-supervised comparison process extracted from environmental invariants and statistical regularities from the stimulus input [75-77]. Future research should investigate which language model can better account for the temporal variations found in speech.

A natural feature of our model is that time can act as a cue for content implemented as a phase code [43, 66]. This code unravels as an ordered list of predictability strength of the internal model. We predict that if speech nodes have a different base activity, ambiguous stimulus interpretation should dependent on the time/phase of presentation (see [31, 63]). Indeed, we could model two temporal speech illusions (Figure 7). There have also been null results regarding the influence of phase on ambiguous stimulus interpretation [47, 78]. For the speech rate effect, when modifying the time of presentation with a neutral entrainer (summed sinusoidals with random phase), no obvious phase effect was reported [47]. A second null result relates to a study where participants were specifically instructed to maintain a specific perception in different blocks which likely increases the pre-activation and thereby the phase [78]. Future studies need to investigate the use of temporal/phase codes to disambiguate speech input and specifically use predictions in their design.

The temporal dynamics of speech signals needs to be integrated with the temporal dynamics of brain signals. However, it is unnecessary (and unlikely) that the exact duration of speech matches with the exact duration of brain processes. Temporal expansion or squeezing of stimulus inputs occur regularly in the brain [79, 80] and this temporal morphing also maps to duration [81-83] or order illusions [84]. Our model predicts increased rhythmic responses for non-isochronous speech matching the internal model. The perceived rhythmicity of speech could therefore also be an illusion generated by a rhythmic brain signal somewhere in the brain.

When investigating the pseudo-rhythmicity in speech it is important to identify situations where speech is actually more rhythmic. Two examples are the production of lists [85] and infant-directed speech [86]. In both these examples it is clear that a strong internal predictive language model is lacking either on the producer’s or on the receiver’s side, respectively. The infant-directed speech also illustrates that a producer might proactively adapt its speech rhythm to the expectations of the internal model of the receiver to align better with the predictions from the receiver’s model (Figure 8B; similar to when you are speaking to somebody that is just learning a new language). Other examples in which speech is more isochronous is during poems, during emotional conversation [87], and in noisy situations [88]. While speculative, it is conceivable that in these circumstances one puts more weight on a different level of hierarchy than the internal linguistic model. In the case of poems and emotional conversation an emotional route might get more weight in processing. In the case of noisy situations, stimulus input has to pass the first hierarchical level of the primary auditory cortex which effectively gets more weight than the internal model.

**Figure 8.**
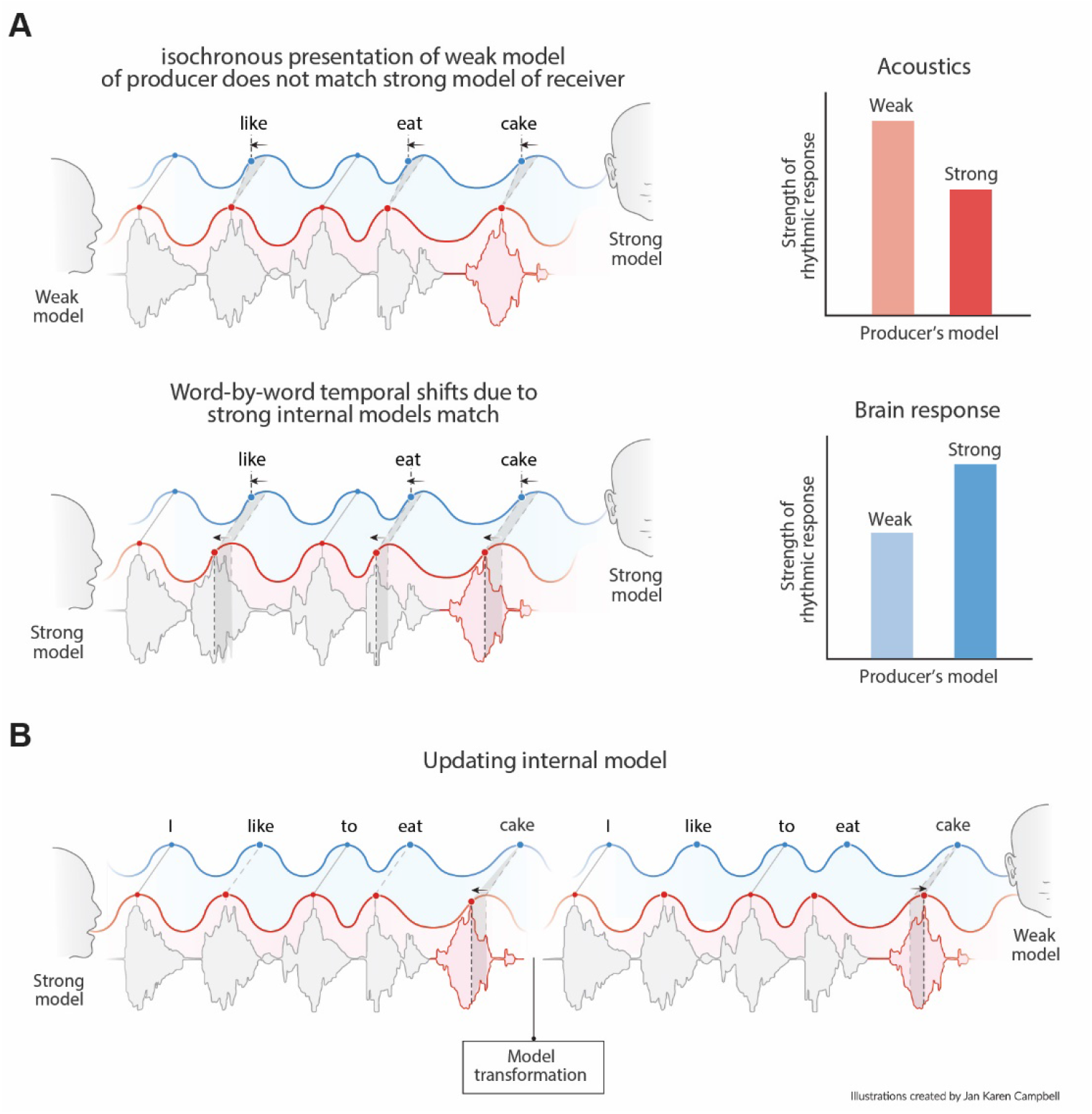
Predictions of the model. A) Acoustics signals will be more rhythmic when a producer has a weak versus a strong internal model (top right). When the producer’s strong model matches the receiver’s model the brain response will be more rhythmic for less rhythmic acoustic input. B) When a producer realizes the model of the receiver is weak it might transform its model and thereby their speech timing to match the receiver’s expectations.

## Conclusions

We argued that pseudo-rhythmicity in speech is in part a consequence of top-down predictions flowing from an internal model of language. This pseudo-rhythmicity is created by a speaker and expected by a receiver if they have overlapping internal language models. Oscillatory tracking of this signal does not need to be hampered by the pseudo-rhythmicity, but can use temporal variations as a cue to extract content information since the phase of activation parametrically relates to the likelihood of an input relative to the internal model. Brain responses can even be more rhythmic to pseudo-rhythmic compared to isochronous speech if they follow the temporal delays imposed by the internal model. This account provides various testable predictions which we list in Table 3 and Figure 8. We believe that by integrating neuroscientific explanations of speech tracking with linguistic models of language processing [21, 24], we can improve to explain temporal speech dynamics. This will ultimately aid our understanding of language in the brain and provide a means to improve temporal properties in speech synthesis.

**Table 3.** Predictions from the current model.

|  |
| --- |
| The more predictable a word, the earlier this word is uttered. |
| When there is a flat constraint distribution over an utterance (e.g., when probabilities are uniform over the utterance) the acoustics of speech should naturally be more rhythmic (Figure 8A). |
| If speech timing matches the internal language model, brain responses should be more rhythmic even if the acoustics are not (Figure 8A). |
| The more similar the internal language models of two speakers, the more effective they are in 'entraining' each other's brain. |
| If speakers suspect their listener to have a flatter constraint distribution than themselves (e.g., the environment is noisy, or the speakers are in a second language context), they adjust to the distribution by speaking more rhythmically (Figure 8B). |
| One adjusts the weight of the constraint distribution to a hierarchical level when needed. For example, when there is noise, participants adjust to the rhythm of primary auditory cortex instead of higher order language models. As a consequence, they speak more rhythmically. |

The theoretical account provides various predictions that are listed in this table.

## Acknowledgments

AEM was supported by the Lise Meitner Research Group “Language and Computation in Neural Systems” from the Max Planck Society, and by the Netherlands Organization for Scientific Research (grant 016.Vidi.188.029). Figure 1 and 8 were created in collaboration with scientific illustrator Jan-Karen Campbell (www.jankaren.com).

## Supporting Figures

**Supporting figure 1.**
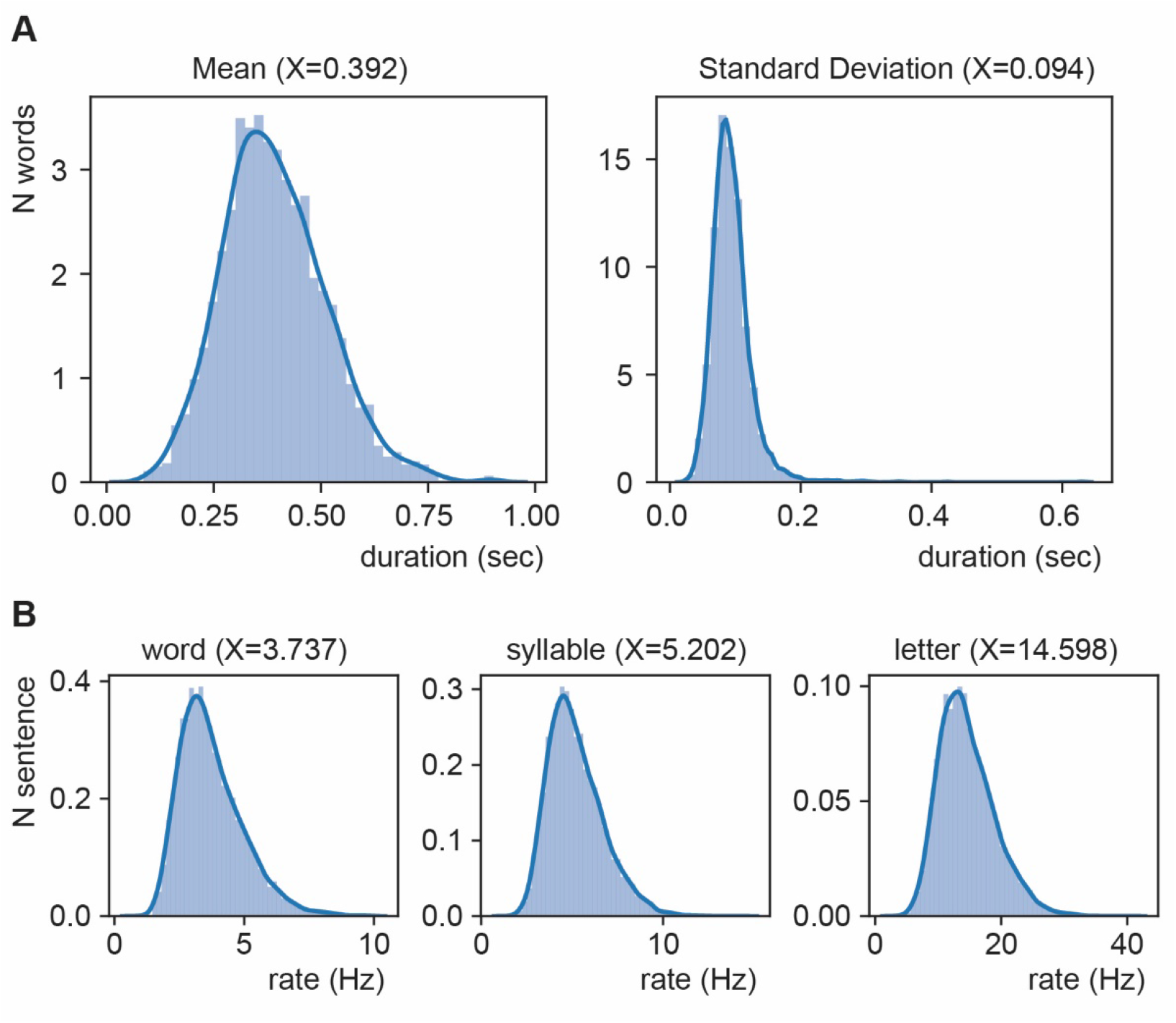
Distribution of mean duration (A) and of average rate (B).

**Supporting figure 2.**
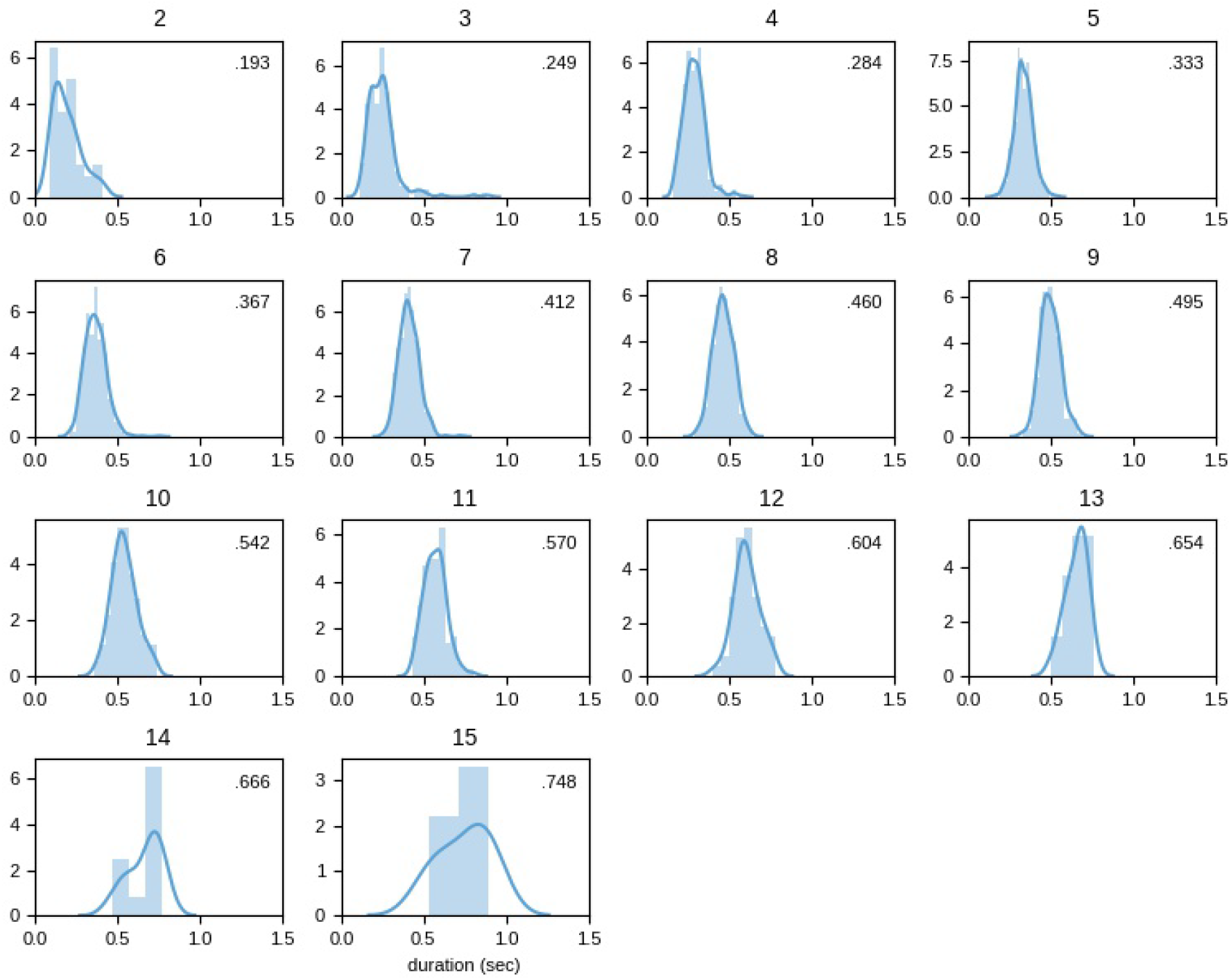
Distribution of mean duration split up for word length (in characters).

**Supporting figure 3.**
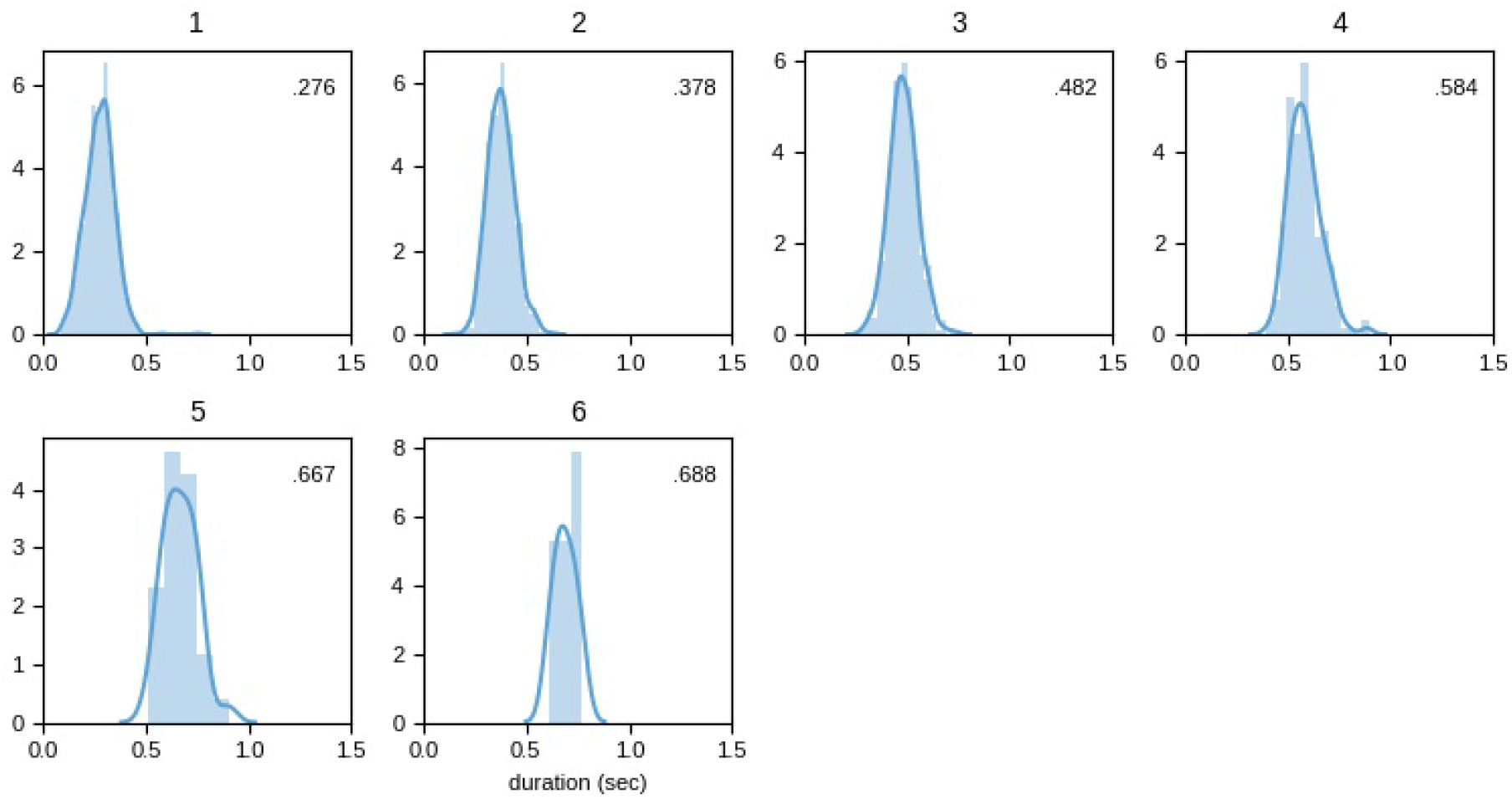
Distribution of mean duration split up for syllable length.

**Supporting figure 4.**
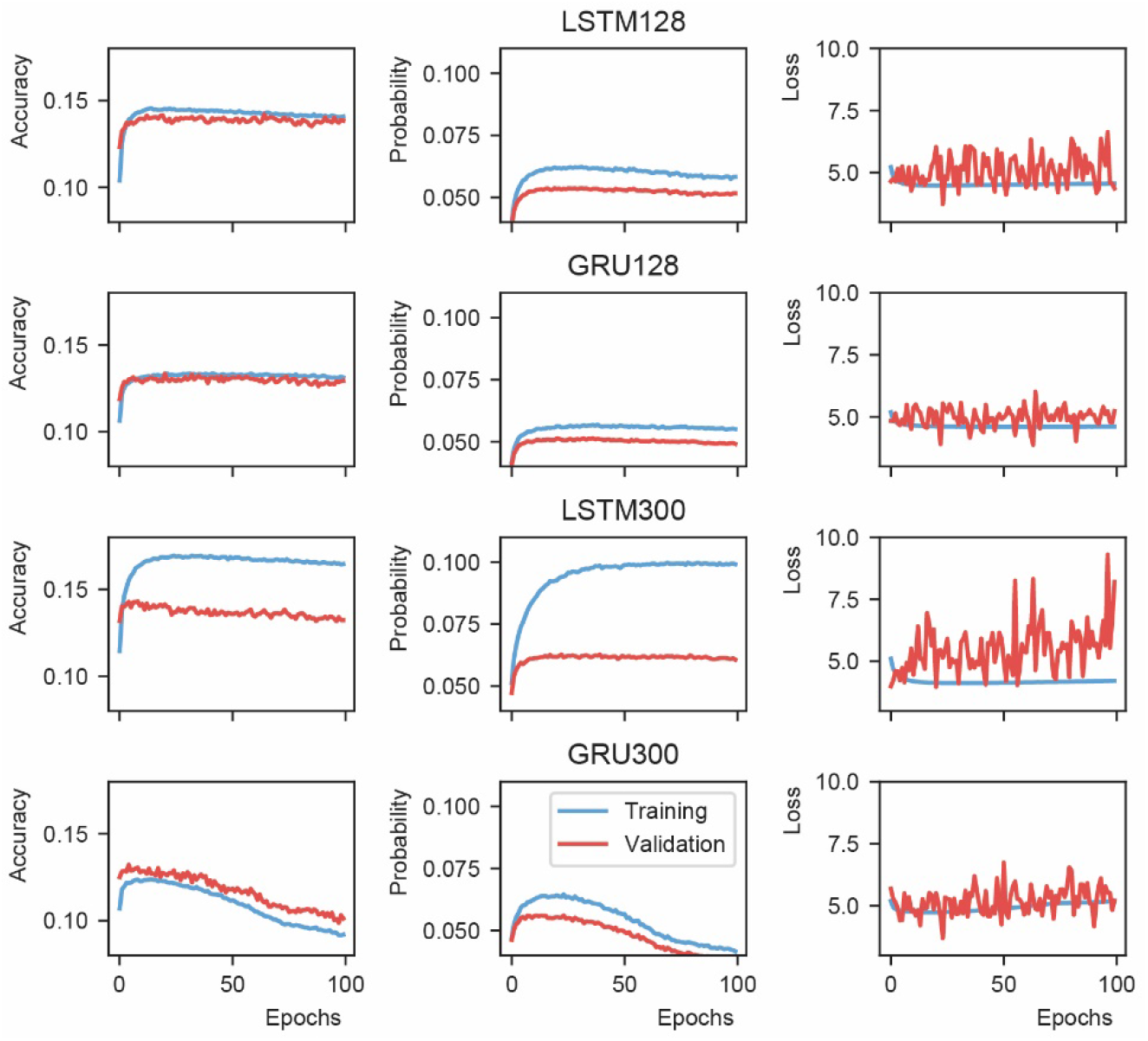
Recurrent neural network evaluation. Probability is defined as the mean of the model output value at the node representing the next word.

**Supporting figure 5.**
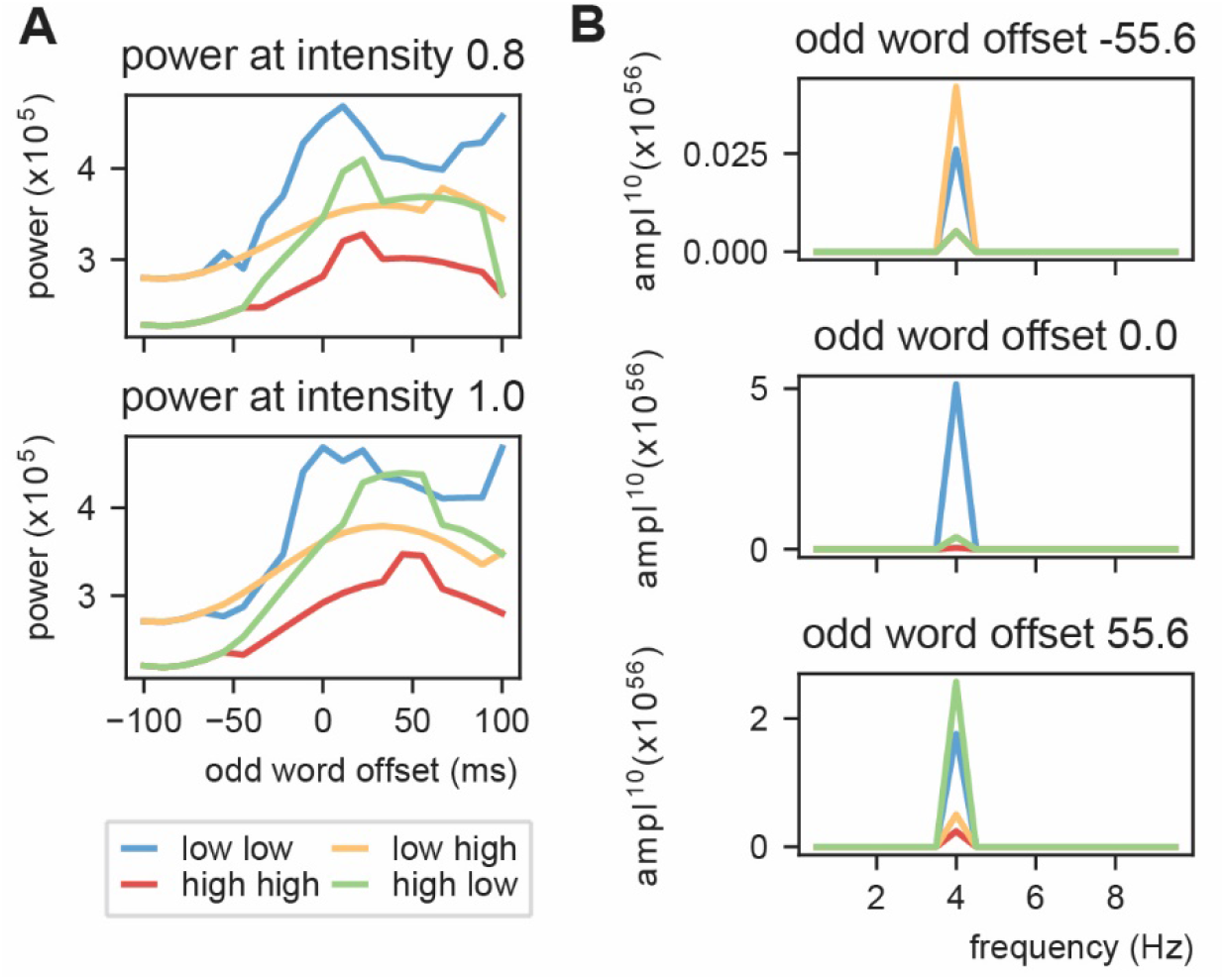
Power at 4 Hz using linearly increasing sensory input. Conventions are the same as in Figure 5D and E.

**Supporting figure 6.**
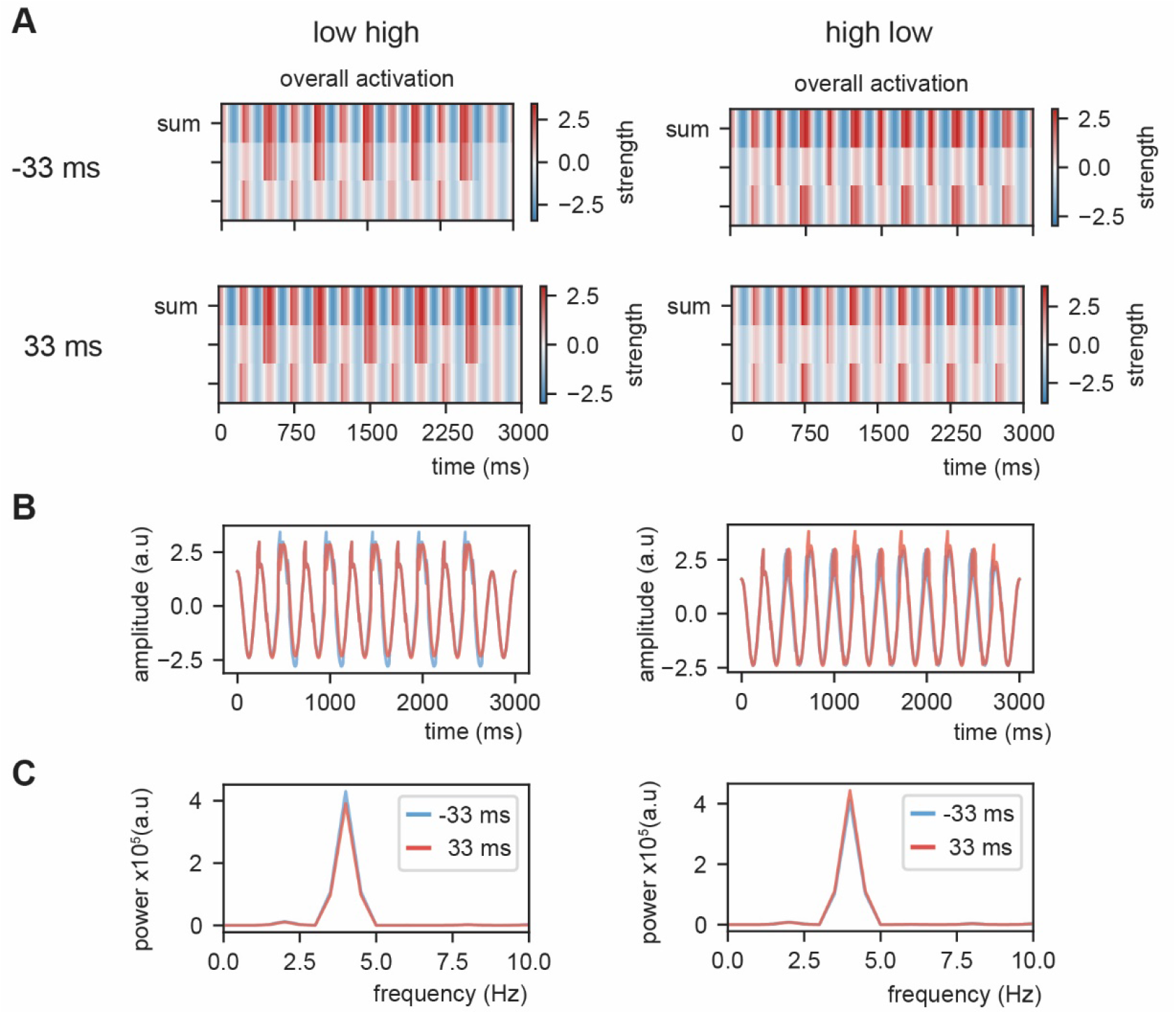
Example of overall activation at threshold 0.8 (gaussian shaped input).

**Supporting figure 7.**
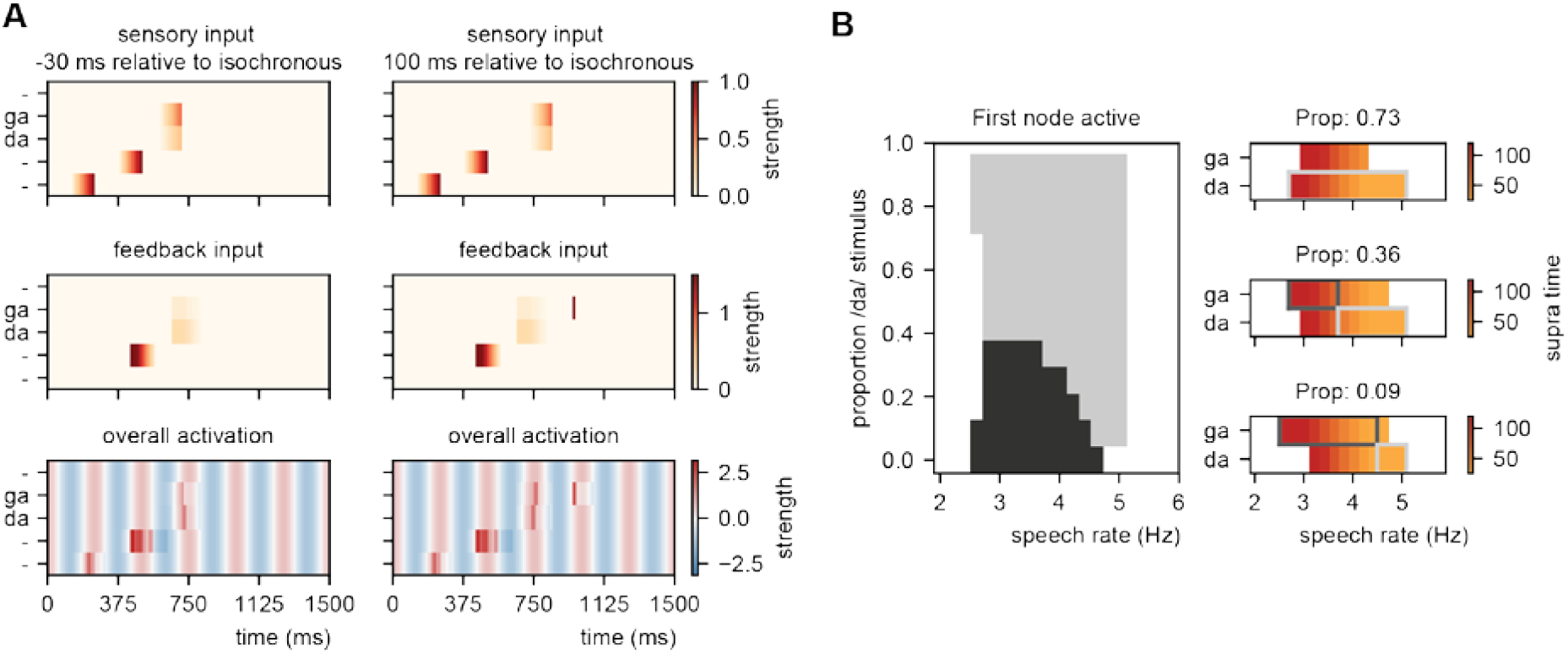
Explaining speech timing illusions. A) Model activation of two example delays for the fitting (figure 7A). B) Modulations due to ambiguous input at different speech rates. Illustration of the node that is active first. Different proportions of the /da/ stimulus show activation timing modulations at different speech rates. Conventions are the same as figure 7A.

## References

1. Jones MR, Boltz M. Dynamic attending and responses to time. Psychological Review. 1989;96(3):459.

2. Large EW, Jones MR. The dynamics of attending: How people track time-varying events. Psychological Review. 1999;106(1):119.

3. Giraud AL, Poeppel D. Cortical oscillations and speech processing: emerging computational principles and operations. Nat Neurosci. 2012;15(4):511–7.

4. Ghitza O, Greenberg S. On the possible role of brain rhythms in speech perception: intelligibility of time-compressed speech with periodic and aperiodic insertions of silence. Phonetica. 2009;66(1-2):113–26.

5. Ding N, Patel AD, Chen L, Butler H, Luo C, Poeppel D. Temporal modulations in speech and music. Neurosci Biobehav Rev. 2017;81:181–7.

6. Arvaniti A. Rhythm, timing and the timing of rhythm. Phonetica. 2009;66(1-2):46–63.

7. Poeppel D. The analysis of speech in different temporal integration windows: cerebral lateralization as ‘asymmetric sampling in time’. Speech Communication. 2003;41(1):245–55.

8. Schroeder CE, Lakatos P. Low-frequency neuronal oscillations as instruments of sensory selection. Trends Neurosci. 2009;32(1):9–18.

9. Luo H, Tian X, Song K, Zhou K, Poeppel D. Neural response phase tracks how listeners learn new acoustic representations. Curr Biol. 2013;23(11):968–74.

10. Keitel A, Gross J, Kayser C. Perceptually relevant speech tracking in auditory and motor cortex reflects distinct linguistic features. PLoS Biol. 2018;16(3):e2004473.

11. Lakatos P, Karmos G, Mehta AD, Ulbert I, Schroeder CE. Entrainment of neuronal oscillations as a mechanism of attentional selection. science. 2008;320(5872):110–3.

12. Henry MJ, Obleser J. Frequency modulation entrains slow neural oscillations and optimizes human listening behavior. Proc Natl Acad Sci. 2012;109(49):20095–100.

13. Herrmann B, Henry MJ, Grigutsch M, Obleser J. Oscillatory phase dynamics in neural entrainment underpin illusory percepts of time. J Neurosci. 2013;33(40):15799–809.

14. Obleser J, Kayser C. Neural entrainment and attentional selection in the listening brain. Trends Cogn Sci. 2019;23(11):913–26.

15. Rimmele JM, Morillon B, Poeppel D, Arnal LH. Proactive sensing of periodic and aperiodic auditory patterns. Trends Cogn Sci. 2018;22(10):870–82.

16. Nolan F, Jeon H-S. Speech rhythm: a metaphor? Philosophical Transactions of the Royal Society B: Biological Sciences. 2014;369(1658):20130396.

17. Jadoul Y, Ravignani A, Thompson B, Filippi P, de Boer B. Seeking temporal predictability in speech: comparing statistical approaches on 18 world languages. Front Hum Neurosci. 2016;10:586.

18. Meyer L. The neural oscillations of speech processing and language comprehension: state of the art and emerging mechanisms. Eur J Neurosci. 2018;48(7):2609–21.

19. Poeppel D, Assaneo MF. Speech rhythms and their neural foundations. Nature Reviews Neuroscience. 2020:1–13.

20. Ten Oever S, Sack AT, Wheat KL, Bien N, Van Atteveldt N. Audio-visual onset differences are used to determine syllable identity for ambiguous audio-visual stimulus pairs. Frontiers in Psychology. 2013;4.

21. Martin AE. Language processing as cue integration: Grounding the psychology of language in perception and neurophysiology. Frontiers in psychology. 2016;7:120.

22. Rosen S. Temporal information in speech: acoustic, auditory and linguistic aspects. Philosophical Transactions of the Royal Society of London Series B: Biological Sciences. 1992;336(1278):367–73.

23. van de Ven V, Kochs S, Smulders F, De Weerd P. Learned interval time facilitates associate memory retrieval. Learn Memory. 2017;24(4):158–61.

24. Martin AE. A compositional neural architecture for language. J Cognit Neurosci. 2020:1–20.

25. cc Marslen-Wilson WD. Functional parallelism in spoken word-recognition. Cognition. 1987;25(1-2):71–102.

26. Lau EF, Phillips C, Poeppel D. A cortical network for semantics:(de) constructing the N400. Nature Reviews Neuroscience. 2008;9(12):920–33.

27. Nieuwland MS. Do ‘early’brain responses reveal word form prediction during language comprehension? A critical review. Neurosci Biobehav Rev. 2019;96:367–400.

28. O’Keefe J, Recce ML. Phase relationship between hippocampal place units and the EEG theta rhythm. Hippocampus. 1993;3(3):317–30.

29. Malhotra S, Cross RW, van der Meer MA. Theta phase precession beyond the hippocampus. Reviews in the neurosciences. 2012;23(1):39–65.

30. Bahramisharif A, Jensen O, Jacobs J, Lisman J. Serial representation of items during working memory maintenance at letter-selective cortical sites. PLoS Biol. 2018;16(8):e2003805.

31. Ten Oever S, Sack AT. Oscillatory phase shapes syllable perception. Proc Natl Acad Sci. 2015;112(52):15833–7.

32. Kayser SJ, McNair SW, Kayser C. Prestimulus influences on auditory perception from sensory representations and decision processes. Proc Natl Acad Sci. 2016;113(17):4842–7.

33. Di Liberto GM, O’Sullivan JA, Lalor EC. Low-frequency cortical entrainment to speech reflects phoneme-level processing. Curr Biol. 2015;25(19):2457–65.

34. Ten Oever S, Hausfeld L, Correia J, Van Atteveldt N, Formisano E, Sack A. A 7T fMRI study investigating the influence of oscillatory phase on syllable representations. NeuroImage. 2016;141:1–9.

35. Thézé R, Giraud A-L, Mégevand P. The phase of cortical oscillations determines the perceptual fate of visual cues in naturalistic audiovisual speech. Science advances. 2020;6(45):eabc6348.

36. Brennan JR, Martin AE. Phase synchronization varies systematically with linguistic structure composition. Philosophical Transactions of the Royal Society B. 2020;375(1791):20190305.

37. Kaufeld G, Bosker HR, Alday PM, Meyer AS, Martin AE. Linguistic structure and meaning organize neural oscillations into a content-specific hierarchy. BioRxiv. 2020.

38. Ghitza O. The theta-syllable: a unit of speech information defined by cortical function. Frontiers in psychology. 2013;4:138.

39. Ghitza O. On the role of theta-driven syllabic parsing in decoding speech: intelligibility of speech with a manipulated modulation spectrum. Frontiers in Psychology. 2012;3.

40. Panzeri S, Macke JH, Gross J, Kayser C. Neural population coding: combining insights from microscopic and mass signals. Trends Cogn Sci. 2015;19(3):162–72.

41. Kayser C, Montemurro MA, Logothetis NK, Panzeri S. Spike-phase coding boosts and stabilizes information carried by spatial and temporal spike patterns. Neuron. 2009;61(4):597–608.

42. Mehta M, Lee A, Wilson M. Role of experience and oscillations in transforming a rate code into a temporal code. Nature. 2002;417(6890):741–6.

43. Lisman JE, Jensen O. The theta-gamma neural code. Neuron. 2013;77(6):1002–16.

44. Reinisch E, Sjerps MJ. The uptake of spectral and temporal cues in vowel perception is rapidly influenced by context. Journal of Phonetics. 2013;41(2):101–16.

45. Kösem A, Bosker HR, Takashima A, Meyer A, Jensen O, Hagoort P. Neural entrainment determines the words we hear. Curr Biol. 2018;28(18):2867-75. e3.

46. Bosker HR, Reinisch E, editors. Normalization for speechrate in native and nonnative speech. 18th International Congress of Phonetic Sciences (ICPhS 2015); 2015: International Phonetic Association.

47. Bosker HR, Kösem A, editors. An entrained rhythm’s frequency, not phase, influences temporal sampling of speech. Interspeech 2017; 2017.

48. O’Malley S, Besner D. Reading aloud: Qualitative differences in the relation between stimulus quality and word frequency as a function of context. Journal of Experimental Psychology: Learning, Memory, and Cognition. 2008;34(6):1400.

49. Monsell S. The nature and locus of word frequency effects in reading. 1991.

50. Monsell S, Doyle MC, Haggard PN. Effects of frequency on visual word recognition tasks: Where are they? Journal of Experimental Psychology: General. 1989;118(1):43.

51. Powers DM, editor Applications and explanations of Zipf’s law. New methods in language processing and computational natural language learning; 1998.

52. Piantadosi ST. Zipf’s word frequency law in natural language: A critical review and future directions. Psychonomic bulletin & review. 2014;21(5):1112–30.

53. Hagoort P. The core and beyond in the language-ready brain. Neurosci Biobehav Rev. 2017;81:194–204.

54. Beattie GW, Butterworth BL. Contextual probability and word frequency as determinants of pauses and errors in spontaneous speech. Language and speech. 1979;22(3):201–11.

55. Gwilliams L, King J-R, Marantz A, Poeppel D. Neural dynamics of phoneme sequencing in real speech jointly encode order and invariant content. bioRxiv. 2020.

56. Deacon D, Mehta A, Tinsley C, Nousak JM. Variation in the latencies and amplitudes of N400 and NA as a function of semantic priming. Psychophysiology. 1995;32(6):560–70.

57. Aubanel V, Schwartz J-L. The role of isochrony in speech perception in noise. Scientific reports. 2020;10(1):1–12.

58. Pellegrino F, Coupé C, Marsico E. A cross-language perspective on speech information rate. Language. 2011:539–58.

59. Thompson SP, Newport EL. Statistical learning of syntax: The role of transitional probability. Language learning and development. 2007;3(1):1–42.

60. Guest O, Martin AE. How computational modeling can force theory building in psychological science. Perspectives on Psychological Science. in press.

61. Meyer L, Sun Y, Martin AE. Synchronous, but not entrained: Exogenous and endogenous cortical rhythms of speech and language processing. Language, Cognition and Neuroscience. 2019:1–11.

62. Meyer L, Sun Y, Martin AE. “Entraining” to speech, generating language? Language, Cognition and Neuroscience. in press.

63. Ten Oever S, Meierdierks T, Duecker F, De Graaf TA, Sack AT. Phase-coded oscillatory ordering promotes the separation of closely matched representations to optimize perceptual discrimination. iScience. 2020:101282.

64. Peelle JE, Davis MH. Neural oscillations carry speech rhythm through to comprehension. Frontiers in Psychology. 2012;3.

65. Martin AE, Doumas LA. A mechanism for the cortical computation of hierarchical linguistic structure. PLoS Biol. 2017;15(3):e2000663.

66. Jensen O, Bonnefond M, VanRullen R. An oscillatory mechanism for prioritizing salient unattended stimuli. Trends in cognitive sciences. 2012;16(4):200–6.

67. Buzsáki G, Draguhn A. Neuronal oscillations in cortical networks. science. 2004;304(5679):1926–9.

68. Bastos AM, Usrey WM, Adams RA, Mangun GR, Fries P, Friston KJ. Canonical microcircuits for predictive coding. Neuron. 2012;76(4):695–711.

69. Michalareas G, Vezoli J, Van Pelt S, Schoffelen J-M, Kennedy H, Fries P. Alpha-beta and gamma rhythms subserve feedback and feedforward influences among human visual cortical areas. Neuron. 2016;89(2):384–97.

70. Lisman JE. The theta/gamma discrete phase code occuring during the hippocampal phase precession may be a more general brain coding scheme. Hippocampus. 2005;15(7):913–22.

71. Cumin D, Unsworth C. Generalising the Kuramoto model for the study of neuronal synchronisation in the brain. Physica D: Nonlinear Phenomena. 2007;226(2):181–96.

72. Chater MHCN. Connectionist psycholinguistics: Greenwood Publishing Group; 2001.

73. McClelland JL, Elman JL. The TRACE model of speech perception. Cognitive psychology. 1986;18(1):1–86.

74. zzz Friederici AD. The brain basis of language processing: from structure to function. Physiological reviews. 2011;91(4):1357–92.

75. Martin AE, Doumas LA. Predicate learning in neural systems: using oscillations to discover latent structure. Current Opinion in Behavioral Sciences. 2019;29:77–83.

76. Doumas LA, Hummel JE, Sandhofer CM. A theory of the discovery and predication of relational concepts. Psychological review. 2008;115(1):1.

77. Doumas LA, Martin AE. Learning structured representations from experience. Psychology of Learning and Motivation. 69: Elsevier; 2018. p. 165–203.

78. Kösem A, Basirat A, Azizi L, van Wassenhove V. High-frequency neural activity predicts word parsing in ambiguous speech streams. J Neurophysiol. 2016;116(6):2497–512.

79. Eagleman DM, Peter UT, Buonomano D, Janssen P, Nobre AC, Holcombe AO. Time and the brain: how subjective time relates to neural time. J Neurosci. 2005;25(45):10369–71.

80. Pariyadath V, Eagleman D. The effect of predictability on subjective duration. PloS one. 2007;2(11):e1264.

81. Eagleman DM. Human time perception and its illusions. Curr Opin Neurobiol. 2008;18(2):131–6.

82. Terao M, Watanabe J, Yagi A, Nishida Sy. Reduction of stimulus visibility compresses apparent time intervals. Nat Neurosci. 2008;11(5):541–2.

83. Ulrich R, Nitschke J, Rammsayer T. Perceived duration of expected and unexpected stimuli. Psychological research. 2006;70(2):77–87.

84. Vroomen J, Keetels M. Perception of intersensory synchrony: A tutorial review. Attention, Perception, & Psychophysics. 2010;72(4):871–84.

85. Jefferson G. List construction as a task and resource. Interaction competence. 1990;63:92.

86. Fernald A. Speech to infants as hyperspeech: Knowledge-driven processes in early word recognition. Phonetica. 2000;57(2-4):242–54.

87. Hawkins S. Situational influences on rhythmicity in speech, music, and their interaction. Philosophical Transactions of the Royal Society B: Biological Sciences. 2014;369(1658):20130398.

88. Bosker HR, Cooke M. Talkers produce more pronounced amplitude modulations when speaking in noise. The Journal of the Acoustical Society of America. 2018;143(2):EL121–EL6.

